# FMNL1 polymerized actin filaments complete mitochondrial division downstream of Golgi-derived vesicles

**DOI:** 10.64898/2026.10.21.760700

**Authors:** Brandon Tabori, Amrapali Ghosh, Maite R. Zavala, Rajarshi Chakrabarti

## Abstract

Mitochondrial division includes contributions from several organelles yet the molecular machinery that facilitates the final Drp1-induced scission of the OMM is largely elusive. We elucidate that FMNL1, a formin family actin assembly factor, is recruited to the mitochondrial division sites through the TGN vesicles in a calcium dependent pathway to complete mitochondrial division. FMNL1 is activated specifically by Cdc42 that generates actin filaments to complete division of the OMM in Drp1 and actin-dependent pathway. Depletion of FMNL1 enhances early events in mitochondrial division pathway characterized by enhanced ER-mitochondrial contacts and increased Drp1 and TGN vesicles on mitochondria which still results in failed mitochondrial division.

## Introduction

Mitochondria are highly dynamic organelles that can modify their morphological network adapting to cellular needs. Mitochondrial dynamics involves division of an expanded network to form smaller units, fusion of smaller units to form elongated network (Chen et al., 2023) and short/long range transport using the microtubules (Shen et al., 2018) and actin network (Moore et al., 2021). Dysregulation of mitochondrial dynamics has been linked to various diseases such as neurodegeneration, cancers, and diabetes (Ma et al., 2020; Cho et al., 2010; Yoon et al., 2011)

Mitochondrial division is a highly orchestrated process the understanding of which have evolved over the years. Mitochondrial division is regulated by the large GTPase dynamin-related protein 1 (Drp1), binding to mitochondrial-bound receptors Mff, MiD49/51, and Fis1. Drp1 then oligomerizes into a ring-like helical structure around a dividing mitochondrion before undergoing a conformational change through GTP hydrolysis that results in its ability to constrict membranes (Kamerkar et al., 2025; Lackner and Nunnari, 2009; Smirnova et al., 2001). In addition to molecular factors, several organelles have been shown to regulate mitochondrial division. The endoplasmic reticulum wraps around the mitochondria and marks the site of mitochondrial division (Friedman et al., 2011) and mtDNA replication prior to the division (Lewis et al., 2016). Mitochondrial division sites have been shown to be positive for lysosomes and are controlled by Rab7 GTP hydrolysis (Wong et al., 2018). More recently, PI3-P laden endosomes have been shown to regulate mitochondrial division in Caenorhabditis elegans (Zhao et al., 2026). Finally, PI4-P positive vesicles from the trans-Golgi network (TGN) have been shown to catalyze the final steps of mitochondrial division in a Drp1 dependent manner (Nagashima et al., 2020). While several organelles are recruited at the mitochondrial division site, the exact mechanism that leads to the final scission of the mitochondria remains elusive.

Actin filaments regulate various aspects of mitochondrial biology including division (Kamerkar et al., 2025), distribution (Moore et al., 2021) and mitochondrial dysfunction induced signaling (Chakrabarti et al., 2022; Fung et al., 2025). INF2 polymerized actin filaments stimulate ER calcium release (Zavala et al., 2026), ER-mitochondrial (ERMC) formation (Chakrabarti et al., 2018) and mitochondrial calcium uptake (Chakrabarti et al., 2018) leading to constriction and division of inner mitochondrial membrane (IMM). INF2-mediated actin filaments also leads to productive Drp1 recruitment onto outer mitochondrial membrane (OMM) thereby stimulating its division (Ji et al., 2015, 2017; Korobova et al., 2013). Additionally, actin filaments can also stimulate Drp1 GTPase activity thereby promoting Drp1 generated constrictive force to divide OMM (Hatch et al., 2016). While INF2 dependent actin assembly occurs in seconds, this actin also depolymerizes quickly making INF2-induced actin turn over fairly short-lived (Chakrabarti et al., 2018; Ji et al., 2015). While Drp1 translocation and oligomerization on mitochondria depends on INF2-mediated actin assembly, yet it occurs kinetically slower than INF2-mediated actin turnover (Chakrabarti et al., 2018; Ji et al., 2015). This required the need to elucidate the real identity of the actin pool that promotes the Drp1 GTP hydrolysis downstream of TGN vesicle recruitment to promote OMM division.

FMNL1 (also known as FRL1) is a bonafide member of the “Formin-Like” family of diaphanous-related formins comprised of FMNL2 and FMNL3 (Schönichen and Geyer, 2010). Similar to other formins, FMNL1 is also auto-inhibited by DID-DAD binding which is relieved by binding of GTPase at the N-terminally located GTPase binding domain (GBD) (Schönichen and Geyer, 2010). FMNL1 has been implicated in podosome dynamics (Mersich et al., 2010), cell migration, phagocytosis, cell adhesion(Yayoshi-Yamamoto et al., 2000; Seth et al., 2006), MTOC regulation (Gomez et al., 2007) and maintenance of Golgi architecture (Colón-Franco et al., 2011). In the present study we discover that actin filaments polymerized by formin FMNL1 catalyze the final steps of mitochondrial division. Depletion of FMNL1 (and not FMNL2 or 3) significantly elongates mitochondrial network in HeLa and U2OS cells and makes them highly resistant to stimulus induced mitochondrial division characterized by lowering of mitochondrial division rates. OMM division in FMNL1 depleted cells occur significantly slower than the matrix separation indicating its role in the final stages of mitochondrial division. Both acute and chronic loss of FMNL1 accentuates the early phases of mitochondrial division characterized by increased ERMC, mitochondrial calcium uptake, Drp1 and TGN translocation to the OMM. Kinetic mapping from live cell images revealed FMNL1 is recruited to mitochondrial division site through TGN vesicles after Drp1 recruitment and before actin filament assembly. In fact, the few fission events in the FMNL1 knockdown cells do not assemble actin filaments during OMM division.

## Results

### FMNL1 depletion induces mitochondrial elongation

To investigate the role of FMNL formins on mitochondrial dynamics, we analyzed mitochondrial morphology in HeLa cells following si-RNA mediated acute depletion of FMNL1 and FMNL2 (**Fig S1A**). Compared to the control cells, FMNL1 depletion caused significant elongation of mitochondrial network (**Fig 1A, B**). FMNL2 depletion had no measurable effect on mitochondrial morphology (**Fig S1B, C**). Mitochondrial morphology in FMNL1 KD cells phenocopied Drp1 KD HeLa cells (**Fig 1A,B**). Since HeLa cells do not express FMNL3, we investigated the role of FMNL3 on mitochondrial morphology in U2-OS cells. Similar to the results obtained in the HeLa cells, knockdown of FMNL1 but not FMNL3 in U2-OS cells displayed a significant elongation in mitochondrial morphology (**Fig S1D-F**). Therefore, from the two cell lines tested it was evident that FMNL1 specifically had an effect on mitochondrial morphology. Furthermore, while acute treatment of HeLa cells with staurosporine and BAPTA-AM caused significant mitochondrial fragmentation in the control cells, FMNL1 KD cells were significantly resistant to such fragmentation similar to Drp1 KD HeLa cells (**Fig 1A, B**). Next, we investigated the extent of mitochondrial network connectivity in FMNL1 depleted cells using the mDendra sensor targeted to mitochondrial matrix that can be irreversibly photoconverted from a green- to red-emitting state when exposed to violet-blue light (Pham et al., 2012). The rate of the acute loss of the photo-converted signal (initial 40 seconds) from the specific ROI was significantly faster (2.5-fold) in the FMNL1 depleted HeLa cells compared to the control cells (**Fig 1C, D**) suggesting a higher interconnectivity of the mitochondrial network upon loss of FMNL1. Since mitochondria in FMNL1 depleted cells resisted agonist-induced mitochondrial fragmentation, we evaluated mitochondrial division events using live-cell microscopy. In unstimulated conditions, while the control cells showed about 5.3 ± 0.2 fission events per min, the rates in FMNL1 depleted cells dropped to 3.78± 0.2 per min (**Fig 1E**). Similar reduction in spontaneous fission rates was also recorded in U2-OS cells (**Fig 1F**). We then evaluated ionomycin-induced (Chakrabarti et al., 2018; Ji et al., 2017) mitochondrial division rates treatment in HeLa cells. Ionomycin-stimulated control cells showed about 11.3 ±0.9 fission events per min while FMNL1 depleted cells clocked about 6.2 ± 0.4 fission events per min (**Fig 1G**). We further evaluated the relative time of IMM-OMM separation in the limited division events that occurred following ionomycin stimulation in the FMNL1 depleted HeLa cells. While in control cells, OMM divided roughly 4.6 ± 0.6 seconds following matrix separation, OMM in FMNL1 KD cells took 10.4 ± 1.4 seconds to divide after matrix separation (**Fig 1H**). In fact, ionomycin stimulated FMNL1 KD cells showed a clear separation of mitochondrial matrix with intact OMM (**Fig S1G**). Furthermore, there were no significant depletion in the abundance of candidate fission and fusion related factors in the FMNL1 KD cells compared to the control cells (**Fig S1H, I**). These results collectively suggest that FMNL1 depletion significantly reduces mitochondrial division events under both unstimulated and stimulated conditions and also leads to slower OMM separation following matrix division. The elongation of mitochondrial network is not a consequence of altered abundance of mitochondrial fission/fusion factors.

**Figure 1.**
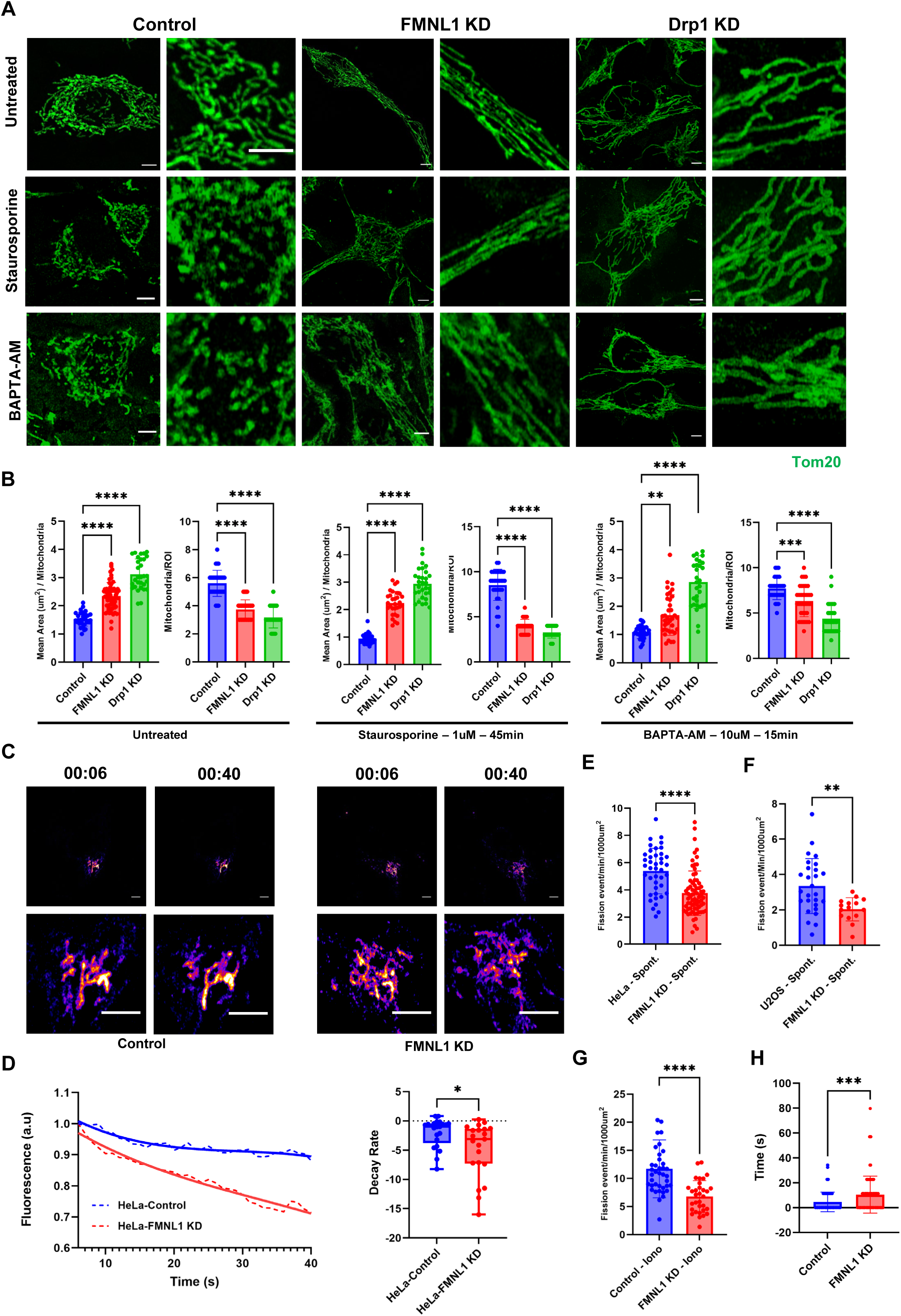
FMNL1 depletion induces mitochondrial elongation. A: Representative fixed-cell images of HeLa Control, FMNL1 KD, and Drp1 KD cells stained for Tom20 (green) before and after treatment with Staurosporine (1µM) for 45min or BAPTA-AM (10µM) for 15min. Scale: 5µm (main and inset) B: Quantification of mitochondrial morphology for each treatment set in (A), quantified for mean area per mitochondria and mean mitochondria per ROI (25µm^2^) as indicated. Data from n = 30 (control), 60 (FMNL1 KD), 30 (Drp1 KD), 31 (Control + staurosporine), 30 (FMNL1 KD + staurosporine), 30 (Drp1 KD + staurosporine), 31 (Control + BAPTA-AM), 36 (FMNL1 KD + BAPTA-AM) and 31 (Drp1 KD + BAPTA-AM) cells from 3 independent experiments. Graphs shown Mean SD. One-way ANOVA used. **** p<0.0001, ** p=0.0011, *** p=0.0008. C: Representative live-cell images of the photoactivated channel of the mito-dendra2 in HeLa Control & FMNL1 KD cells at the time of stimulation (6 seconds) and at 40 seconds. Scale: 5µm. D. Traces representing average decay rate of normalized fluorescence signal at stimulated ROI (25µm^2^) (Left); Plots of decay rate (slopes) for individual cells shown in C. n = 21 for HeLa Control and FMNL1 KD across 3 independent experiments. Unpaired student’s t-test used. * p = 0.0172. E. Quantification of unstimulated mitochondrial division events (event/min/1000µm^2^) in HeLa Control & FMNL1 KD cells transfected with mitoGFP. Data from n = 41 (control) and 71 (FMNL1 KD) cells across 3 – 4 independent experiments Graphs shown Mean SD. Unpaired student’s t-test used. **** p<0.0001 F. Quantification of unstimulated mitochondrial division events (event/min/1000µm^2^) in U2-OS control & FMNL1 KD cells transfected with mitoGFP. Data from n = 21 and 15 cells across 3 independent experiments. Graphs shown Mean SD. Unpaired student’s t-test used. ** p=0.0035 G. Quantification of mitochondrial division events (event/min/1000µm^2^) in HeLa control & FMNL1 KD cells co-transfected with mitoBFP and Tom20-GFP stimulated with ionomycin (5µM). Data from n = 39 and 32 cells across 3 independent experiments for Control and FMNL1 KD, respectively. Graphs shown Mean SD. Unpaired student’s t-test used. **** p<0.0001. H. Quantification of lag-time between IMM and OMM division in HeLa Control & FMNL1 KD cells co-transfected with mitoBFP and Tom20-GFP stimulated with ionomycin (5µM). Data from n = 148 (Control) and 85 (FMNL1 KD) fission events from 15 (Control and FMNL1 KD) cells across 2 independent experiments. Graphs shown Mean SD. Unpaired student’s t-test used. *** p=0.0001.

### FMNL1 depletion potentiates early events in mitochondrial division pathway

We next wanted to evaluate the specific steps in the mitochondrial division pathway perturbed by FMNL1. Ionomycin-induced INF2-mediated actin-burst was unchanged in FMNL1 depleted cells (**Fig S2A, B**) suggesting INF2 activity is not modulated by FMNL1. Next, we measured ER and mitochondrial calcium transients using ERGCamp150 and mito-R-Geco respectively in control and FMNL1 depleted HeLa cells. While histamine stimulated ER calcium release was similar in control and FMNL1 depleted cells (**Fig S2C, D**), FMNL1 KD cells showed a significantly higher mitochondrial calcium uptake (**Fig S2E, F**). This suggested that there might be an increase in ER-Mitochondrial coupling in FMNL1 KD cells leading to increased mitochondrial calcium uptake. To test the effect of FMNL1 KD on ERMC, we first used Proximity Ligation Assay (PLA) against VAP-B (ER) and Tom20 (mitochondria). Interestingly FMNL1 KD showed a significant increase in density and number of PLA dots, suggesting an overall increase in ERMC (**Fig 2A, B**). To further examine this phenotype, we evaluated ERMC from thin-section TEM images. Compared to control cells, FMNL1 KD in HeLa cells significantly increased the extent of close contact between ER and mitochondria (**Fig 2C, D**), a significant decrease in the minimum distance between ER and mitochondria and a significant increase in overall mitochondrial perimeter (**Fig 2E, F**). These results suggest that dampening of mitochondrial division in FMNL1 KD cells is not due to lack of ERMC and/or mitochondrial calcium availability, both of which were significantly strengthened in FMNL1 KD cells.

**Figure 2.**
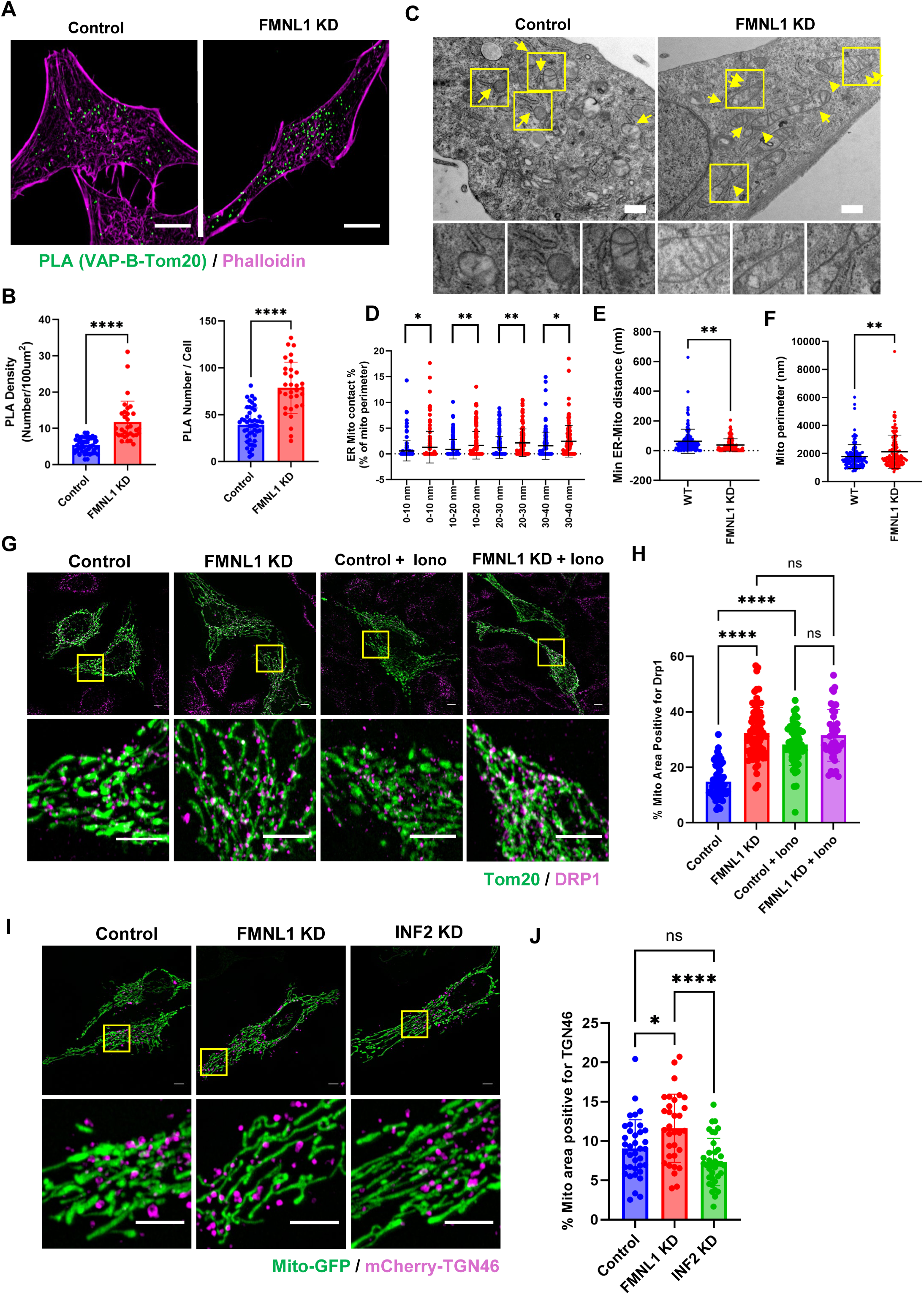
FMNL1 depletion potentiates early events in mitochondrial division. A: Representative fixed-cell images from Proximity Ligation Assay (PLA) against VAP-B and Tom20 (PLA dots in green) in a HeLa Control and FMNL1 KD cell stained for actin filaments with Phalloidin (magenta). Scale: 10µm. B. Quantification of PLA density expressed as number of dots per 100µm^2^ (left) and PLA dots, expressed as number/cell (right). Data from 48 (Control) and 32 (FMNL1 KD) cells from 3 independent experiment. Graph shows Mean ± S.D. Unpaired student’s t-test used. **** p<0.0001. C: Representative images from thin-section TEM in HeLa Control & FMNL1 KD cells showing ER and mitochondria in the sections. The arrows represent close contact between ER and Mitochondria in the sections. Scale: 200 nm. D: Scatter plots showing the percentage of close ER-Mito contacts, expressed as % of mitochondrial perimeter having an ER between 0-10, 10-20, 20-30, and 30-40 nm. Data of 129 (Control) & 124 (FMNL1 KD) mitochondria, from 6 independent experiments. Graphs show Mean ± S.D. Unpaired student’s t-test used. * p=0.0264 (0-10nm), ** p=0.0077 (10-20nm), ** p=0.0021 (20-30nm), *p=0.0167 (30-40nm). E: Scatter plots showing the minimum distance (in nm) between a mitochondrion and an ER. Data of 129 (Control) & 124 (FMNL1 KD) mitochondria, from 6 independent experiments. Graphs show Mean ± S.D. Unpaired student’s t-test used. ** p=0.0029. F: Scatter plots showing perimeter of individual mitochondria (in nm). Data of 129 (Control) & 124 (FMNL1 KD) mitochondria, from 6 independent experiments. Graphs show Mean ± S.D. Unpaired student’s t-test used. ** p=0.0072. G: Representative fixed-cell images of HeLa Control & FMNL1 KD stained for Tom20 (green) and Drp1 (magenta) in unstimulated and ionomycin (5µM/ 5 min) conditions (left). Scale 5µm. H: Quantifications of % mitochondrial area positive for Drp1. Data from n = 53 (Control), 72 (FMNL1 KD), 53 (Control + Iono), and 39 (FMNL1 KD + Iono) cells from 3 independent experiments. Graphs show Mean ± S.D. One-way ANOVA used. **** p<0.0001. I: Representative fixed-cell images of HeLa Control, FMNL1 KD, & INF2 KD cells transfected with mitoGFP (green) and mCherry-TGN46 (magenta). Scale: 5µm. J: Quantifications of % mitochondrial area positive for TGN46. Data from n = 37 (Control), 32 (FMNL1 KD), & 36 (INF2 KD) cells from 3 independent experiments. Graphs show Mean ± S.D. One-way ANOVA used. * p=0.0128, **** p<0.0001.

Next, we evaluated whether Drp1 recruitment onto mitochondria was hampered upon FMNL1 depletion. While 14.9 ± 0.6 % of mitochondrial area was positive for Drp1 in control cells, FMNL1 depleted cells had 32.3 ± 1.0 % of mitochondrial area positive for Drp1 (**Fig 2G, H**). Interestingly, ionomycin induction increased Drp1-positive mitochondrial area only in control cells (28.2 ± 1%) but not in FMNL1 depleted cells (31.5 ± 1.4%) (**Fig 2G, H**). Similar results were also seen in U2-OS cells (**Fig S2G, H**). Interestingly while overexpression of Drp1 in control cells caused significant fragmentation of the mitochondrial network, mitochondrial network in FMNL1 KD cells were significantly resistant to fragmentation (**Fig S2I, J**). Therefore, it was evident that similar to ERMCs, Drp1 on mitochondria was also enhanced upon FMNL1 depletion but seemed substantially compromised for mitochondrial division.

Next, we investigated the abundance of TGN vesicles on mitochondria. Control and FMNL1 KD HeLa cells were transfected with mCherry-TGN46 and mitoGFP, fixed and quantified for mitochondrial area positive for TGN46. Compared to control cells, FMNL1 KD cells had significantly higher TGN46 positive mitochondrial area (**Fig 2I, J**). We further compared this to INF2 KD cells. Contrary to the effects of FMNL1 KD, depletion of INF2 had no significant effect on the abundance of TGN vesicles on mitochondria (**Fig 2I, J**). We documented similar results in U2-OS cells as well (**Fig S2K, L**). Thus, loss of FMNL1 potentiates the early events in the mitochondrial division pathway characterized by an increase in ERMC, mitochondrial calcium uptake and mitochondrial recruitment of Drp1 and TGN vesicles.

### Chronic loss of FMNL1 in HeLa cells recapitulates phenotypes of FMNL1 KD cells

To investigate whether loss of FMNL1 induced defects in mitochondrial fission could be compensated over time, we created CRISPR-mediated knockout (KO) of FMNL1 in HeLa cells (**Fig S3A**). Similar to acute loss of FMNL1, FMNL1 KO cells displayed elongated mitochondrial morphology compared to the control cells (**Fig S3B, C**). Live-cell assays showed a significant reduction in rate of unstimulated fission events in FMNL1 KO cells (**Fig S3D**). Further, we stained the FMNL1 KO cells for Drp1 and observed a significant increase in mitochondrial localized Drp1 in unstimulated conditions compared to the control cells (**Fig S3E, F**). Ionomycin stimulation triggered an increase in Drp1 puncta on the mitochondria in control cells but not in FMNL1 KO cells suggesting that chronic depletion of FMNL1 significantly increased mitochondrially localized Drp1 (**Fig S3E, F**). Finally, we also evaluated the extent of TGN vesicles on mitochondria by transfecting in mCherry-TGN46 in control and FMNL1 KO cells.

Similar to results obtained previously, FMNL1 KO cells displayed increased abundance of TGN vesicles on mitochondria (**Fig S3G, H**). Thus, chronic loss of FMNL1 also accentuates early events in the mitochondrial division pathway which is not significantly compensated.

### FMNL1 is recruited to sites of mitochondrial division

Next, we investigated whether FMNL1 is physically present at the mitochondrial division sites or modulates it through a signaling cascade. Transfection of C-terminally tagged FMNL1 (FMNL1-GFP) in HeLa (**S4A-C**) and U2-OS cells (**S4D-F**) displayed a punctate intracellular distribution with increased abundance at the PM. Over-expression of FMNL1-GFP did not cause drastic mitochondrial fragmentation in both cell types (**S4A, B, D, E**). Further analysis revealed an approximately 3.63 ± 0.44 % and 4.19 ± 0.5% overlap of FMNL1-GFP puncta with mitochondria (Tom20) in HeLa (**S4C**) and U2-OS (**S4F**) cells respectively suggesting that a pool of FMNL1 co-localizes with the mitochondria. We also noticed FMNL1-GFP at mitochondrial constriction site together with phalloidin puncta (actin filaments) further suggesting that FMNL1 might be present at mitochondrial division site (**Fig S4G**). To analyze this further we evaluated spontaneous division events for the presence of FMNL1-GFP at mitochondrial division sites in HeLa (**S4H**) and U2-OS cells (**S4I**). Approximately 85% and 81% of spontaneous mitochondrial division events in HeLa and U2-OS cells respectively were positive for the presence of FMNL1-GFP. Therefore, a pool of FMNL1is present on mitochondria and a significant fraction of unstimulated mitochondrial division events were positive for the presence of FMNL1.

Since mitochondrial division proceeds through a concerted effort of Drp1 and TGN46 vesicles, we wanted to co-localize FMNL1 at division sites with respect to the above-mentioned factors. HeLa cells expressing mitoBFP and FMNL1-GFP were transfected with either mCherry-Drp1 (**Fig 3A, D**) or mCherry-TGN46 (**Fig 3B, E**) and evaluated for kinetics of FMNL1 with respect to Drp1 or TGN46 in stimulated (Fig 3A-C) and 4µM ionomycin stimulated (Fig 3D-F) conditions. FMNL1 co-localized with Drp1 (**Fig 3A, D, C, F**) and TGN46 (**Fig 3B, E, C, F**) at both unstimulated and ionomycin-induced mitochondrial division sites. Similar to spontaneous mitochondrial division events, 82% of the division events stimulated by ionomycin were positive for the presence of FMNL1-GFP across all the recordings tested (**S4H)**. In comparison, Drp1 was present in 100% and TGN46 in 72% of both stimulated and unstimulated division events (**Fig S3H**). Further analysis of these division events revealed that FMNL1-GFP arrived at the division site significantly later than mCherry-Drp1 and at a similar time as mCherry-TGN46 for both unstimulated and stimulated mitochondrial division (**Fig 3C, F**). Therefore, a temporal kinetic plot accumulated form all the recorded events revealed that during unstimulated mitochondrial division Drp1 arrived at the division site at approximately 72.8 ± 5.3 seconds prior to a division event, closely followed by TGN46 vesicles (44.9 ± 3 seconds) and FMNL1(44.06 ± 3 seconds) before a division event (**Fig 3C**). The numbers were similar during ionomycin stimulation where Drp1 arrived at the division site at approximately 78.6 ± 4.7 seconds prior to a division event, closely followed by TGN46 vesicles (54.7 ± 3 seconds) and FMNL1(48.0 ± 2 seconds) before a division event.

**Figure 3.**
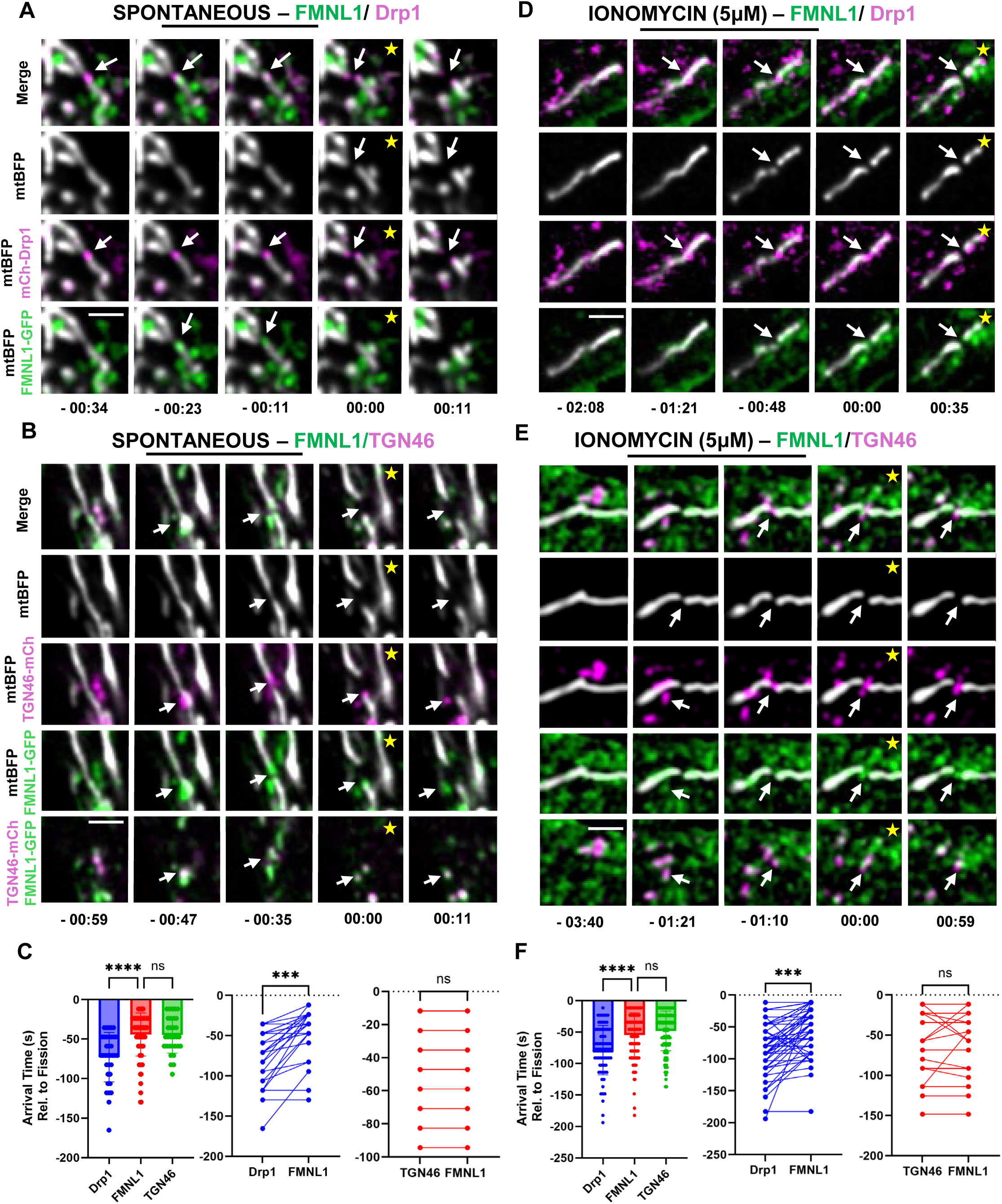
FMNL1 is recruited to mitochondrial fission sites. A: Representative timelapse of an unstimulated mitochondrial division event in HeLa WT cell transfected with mitoBFP (gray), Drp1-mCherry (magenta), and FMNL1-GFP (green). Arrows indicate site of division. Star (yellow) indicate the time-frame of division which is taken as 0 B: Representative timelapse of an unstimulated mitochondrial division event in HeLa WT cell transfected with mitoBFP (gray), mCherry-TGN46 (magenta), and FMNL1-GFP (green). Arrows indicate site of division. Star (yellow) indicate the time-frame of division which is taken as 0 C: (Left) Quantification of arrival times (relative to fission: time 0) of fission factors (Drp1, TGN46, FMNL1). Data from n = 30 (Drp1), 29 (TGN46), 30 (FMNL1) fission events from 10 cells across 2 independent experiments. Graphs show Mean ± S.D. *** p=0.0002. (Right) Quantification of paired arrival time (relative to fission) of Drp1 vs. FMNL1 and TGN46 Vs FMNL1 in the same division event. Graphs show individual points that represent a division event with a connecting line representing points are from same division event. Unpaired student’s t-test used. *** p=0.0002. D: Representative timelapse of a mitochondrial division event in HeLa WT cell transfected with mitoBFP (gray), Drp1-mCherry (magenta), and FMNL1-GFP (green), stimulated with 4µM ionomycin and imaged for 5 minutes. Arrows indicate site of division. Star (yellow) indicate the time-frame of division which is taken as 0 E: Representative timelapse of a mitochondrial division event in HeLa WT cell transfected with mitoBFP (gray), mCherry-TGN46 (magenta), and FMNL1-GFP (green), stimulated with 4µM ionomycin and imaged for 5 minutes. Arrows indicate site of division. Star (yellow) indicate the time-frame of division which is taken as 0 F: (Left) Quantification of arrival times (relative to fission: time 0) of fission factors (Drp1, TGN46, FMNL1). Data from n = 79 (Drp1), 267 (TGN46), 187 (FMNL1) fission events from 15 (Drp1), 61 (TGN46), 37 (FMNL1) cells across 2-4 independent experiments. Graphs show Mean ± S.D. *** p=0.0002. (Right) Quantification of paired arrival time (relative to fission) of Drp1 vs. FMNL1 and TGN46 Vs FMNL1 in the same division event. Data from n = 66 fission events from 15 cells across 3 independent experiments (Drp1 Vs. FMNL1) and Data from n = 33 fission events from 10 cells across 2 independent experiments (FMNL1 Vs TGN46). Graphs show individual points that represent a division event with a connecting line representing points are from same division event. Unpaired student’s t-test used. *** p=0.0001.

While we saw a significant pool of FMNL1 on mitochondria, we observed that following ionomycin stimulation there is significant co-localization of mCherry-TGN46 vesicles with FMNL1-GFP at the division site (**Fig 3B, E; S4J-L**). Indeed, ionomycin treatment stimulated the recruitment of FMNL1-GFP on the mCherry-TGN46 vesicles (**Fig S4J-L**) suggesting that the division-competent pool of FMNL1 might be piggy-backing on TGN46 vesicles to reach mitochondrial division site. We also performed this experiment in the INF2-KO cells which did not dampen ionomycin-induced recruitment of FMNL1 on TGN46 vesicles (**Fig S4J-L**) suggesting that this recruitment is independent of INF2-mediated actin filaments. Overall, our results reveal that a pool of FMNL1 is physically present at mitochondrial division sites being recruited after Drp1 mostly riding on TGN46 vesicles in an INF2-independent manner.

### CDC42 likely activates FMNL1 and is required for mitochondrial fission in a Drp1 dependent manner

FMNL1 contains a GBD domain at its N-terminus and has been shown to be activated either by Cdc42(Chua et al., 2024) or Rac1(Favaro et al., 2010) in a context dependent manner (Kühn and Geyer, 2014). We therefore wanted to uncover the GTPase responsible for activating FMNL1 in the context of mitochondrial division. HeLa cells overexpressing constitutive active Cdc42 (Cdc42-Q61L) showed significant mitochondrial fragmentation compared to the un-transfected control cells (**Fig 4A, E; S5A**). However, overexpression of constitutive active Rac1 (Rac1-Q61L) did not cause mitochondrial fragmentation (**Fig 4A, E; S5A**) in the cells suggesting that activated Cdc42 and not Rac1 is important for mitochondrial division. Next, we wanted to test whether Cdc42 operated in an FMNL1 and/or Drp1 dependent manner to mediate mitochondrial division. Unlike control cells, both FMNL1 KD and Drp1 cells overexpressing either Cdc42-Q61L or Rac1-Q61L did not result in mitochondrial fragmentation (**Fig 4B, C, E**). Thus, it is evident that active Cdc42 regulates mitochondrial division in an FMNL1 and Drp1-dependent manner. Surprisingly, overexpression of Cdc42-Q61L could significantly fragment the elongated mitochondria in INF2-depleted HeLa cells (**Fig 4D, E**) suggesting that significant upregulation of Cdc42 and possibly FMNL1 downstream could somewhat overcome the mitochondrial division blockade induced by the loss of INF2. Furthermore, we wanted to assess whether active Cdc42 localizes to mitochondria. HeLa cells transfected with Cdc42-Q61L showed 24.61±0.8% mitochondrial area positive for active Cdc42 which was independent of FMNL1 (**Fig S5B, S5C**). Interestingly a pool of active Cdc42 also localized to TGN46 vesicles which was slightly dependent on the presence of FMNL1 (**Fig S5D**). However, unlike FMNL1, active Cdc42 was not dynamically recruited to TGN46 vesicles upon stimulation with ionomycin (**Fig S5E**). Live-cell imaging of mitochondrial division events in HeLa cells revealed that active Cdc42 was present at 79% of all the ionomycin-induced mitochondrial division events (**Fig 4F, G**). At these events, roughly 45% of the time Cdc42 was on TGN vesicles while the rest 55% of the time was directly present on mitochondria (**Fig 4H**). We also carried out GFP-trap pull down of FMNL1 in control and active Cdc42 overexpressing cells which revealed an enhanced enrichment in abundance of mitochondrial proteins PTPIP51 and MFF in eluates from cells overexpressing active Cdc42 (**Fig 4I**). This suggests that activation of Cdc42 increases the abundance of FMNL1 on mitochondria, thereby promoting mitochondrial division in an FMNL1 and Drp1 dependent manner.

**Figure 4.**
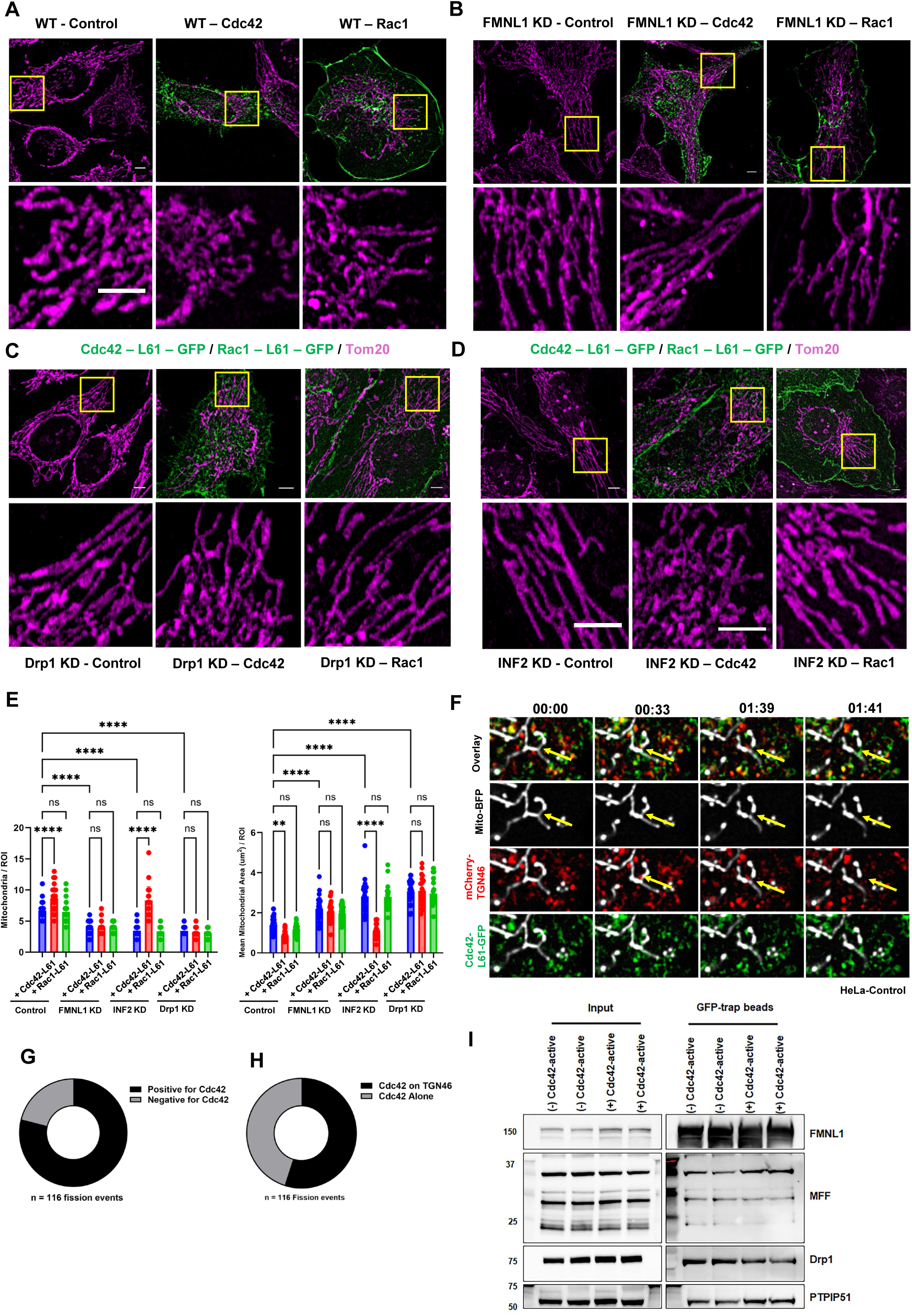
CDC42 likely activates FMNL1 and is required for mitochondrial fission in a Drp1 dependent manner. A: Representative fixed-cell images of HeLa WT cells transfected with either GFP-Cdc42-Q61L (green) or GFP-Rac1-Q61L (green) stained with Tom20 (magenta). Scale: 5µm. B: Representative fixed-cell images of HeLa FMNL1 KD cells transfected with either GFP-Cdc42-Q61L (green) or GFP-Rac1-Q61L (green) stained with Tom20 (magenta). Scale: 5µm. C: Representative fixed-cell images of HeLa Drp1 KD cells transfected with either GFP-Cdc42-Q61L (green) or GFP-Rac1-Q61L (green) stained with Tom20 (magenta). Scale: 5µm. D: Representative fixed-cell images of HeLa INF2 KD cells transfected with either GFP-Cdc42-Q61L (green) or GFP-Rac1-Q61L (green) stained with Tom20 (magenta). Scale: 5µm. E: Quantification of mitochondrial morphology for each set shown in panels A-D. Quantified for mean area per mitochondria (right) and mean mitochondria per ROI (25µm^2^) (left). Data from n = 30 (WT-control), 30 (WT-Cdc42-Q61L), 30 (WT-Rac1-Q61L), 31 (FMNL1 KD-control), 30 (FMNL1 KD-Cdc42-Q61L), 30 (FMNL1 KD-Rac1-Q61L), 30 (Drp1 KD-control), 22 (Drp1 KD-Cdc42-Q61L), 19 (Drp1 KD-Rac1-Q61L), 30 (INF2 KD-control), 30 (INF2 KD-Cdc42-Q61L), 22 (INF2 KD-Rac1-Q61L), from 3 independent experiment. Graphs show Mean ± S.D. One-way ANOVA used. **** p<0.0001, ** p= 0.0014, *** p=0.0002. F: Representative timelapse images of a mitochondrial division event in HeLa WT cell transfected with mitoBFP, mCherry-TGN46, and GFP-Cdc42-Q61L stimulated with 5µM of ionomycin. Arrows indicate site of division. Scale: 5µm. G: Quantification of % division events positive for GFP-Cdc42-Q61L puncta in HeLa WT cells. Data from n = 116 fission events from 25 cells from 2 independent experiment. H: Quantification of % division events GFP-Cdc42-Q61L colocalized with TGN46 prior to division in HeLa WT cells. Data from n = 116 fission events from 25 cells from 2 independent experiment. I: Western blot images of candidate proteins from GFP-trap pull-down assay, in HEK WT cells transfected with FMNL1-GFP with or without Cdc42-G12V (constitutive active). Data Shown from 2 independent pull downs.

### FMNL1 activation partially overrides the loss of ARF1 and PI4-P to mediate mitochondrial division

Arf1 and PI(4)KIIIβ localized TGN-derived vesicles mediate final steps of mitochondrial division through formation of PI4-P at mitochondrial division sites (Nagashima et al., 2020). Indeed, similar to the previous study, we detected a pool of Arf1 and PI4-P co-localized with TGN vesicles (**Fig 5A-D**). While ionomycin stimulation did not enhance Arf1 localization to TGN vesicles (**Fig 5A, C**), we detected a significant increase in PI4-P abundance on TGN vesicles following the stimulation (**Fig 5B, D**). FMNL1 depletion neither reduced Arf1 localization on TGN vesicles (**Fig 5A, C**) nor inhibited ionomycin-induced PI4-P formation on TGN vesicles (**Fig 5B, D**) suggesting that FMNL1 does not regulate either of these factors during mitochondrial division. Reciprocally neither inhibition of Arf1 nor PI(4)KIIIβ altered the dynamics of FMNL1 on TGN upon ionomycin stimulation (**Fig 5E, F**). These results showed that FMNL1 operated independent of Arf1 or PI4-P in mediating mitochondrial division. As previously shown, Arf1 or PI(4)KIIIβ depletion also led to mitochondrial elongation in our system (**Fig 5G, H**). However, over-expression Cdc42-Q61L could significantly fragment mitochondria in both Arf1 and PI(4)KIIIβ depleted cells (**Fig 5G, H**). Taken together these results suggest that while TGN-localized Arf1 and PI(4)KIIIβ are important mediators of mitochondrial division, activating the terminal factor FMNL1 could significantly over-ride the loss of these factors and mediate mitochondrial division in a Drp1-dependent manner.

**Figure 5.**
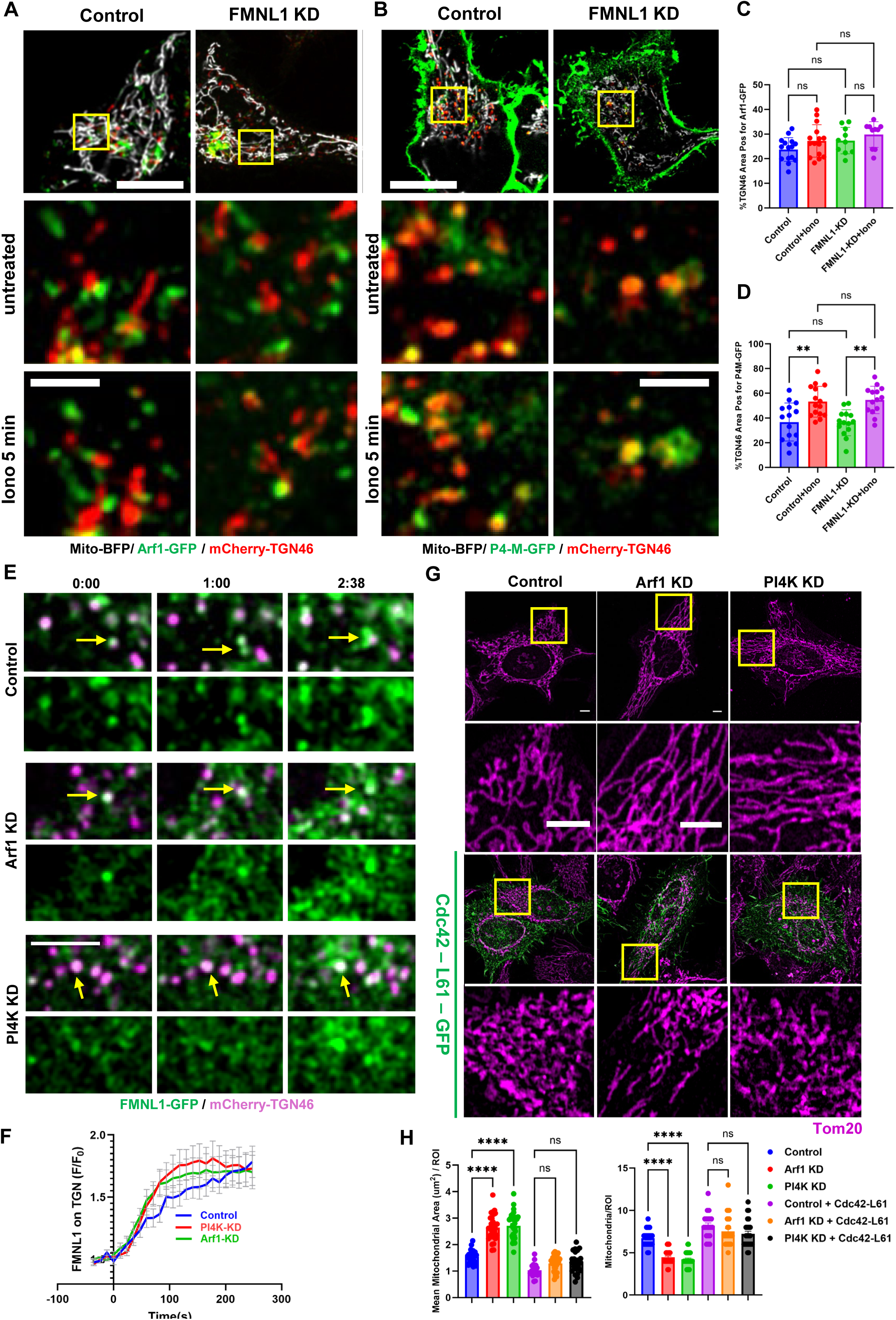
FMNL1 activation partially overrides the loss of ARF1 and PI4-P to mediate mitochondrial division. A: Representative live-cell images from HeLa control & FMNL1 KD cells transfected with mitoBFP (gray), Arf1-GFP (green), and mCherry-TGN46 (red) stimulated with 5µM of ionomycin. Scale: 10µm (main), 5µm (inset). B: Representative live-cell images from HeLa control & FMNL1 KD cells transfected with mitoBFP (gray), P4-M-GFP (green), and mCherry-TGN46 (red) stimulated with 5µM of ionomycin. Scale: 10µm (main), 5µm (inset). Arrows indicate vesicles with increasing P4M-GFP (PI4-P) signal. C: Quantification of %TGN46 area positive for Arf1-GFP in HeLa Control & FMNL1 KD cells shown in (A) before and after stimulation with ionomycin. Data from n = 15 (Control) & 10 (FMNL1 KD) cells from 2 independent experiment. Graphs show Mean ± S.D. One-way ANOVA used. D: Quantification of %TGN46 area positive for P4-M-GFP in HeLa Control & FMNL1 KD cells shown in (B) before and after stimulation with ionomycin. Data from n = 15 (Control) & 14 (FMNL1 KD) cells from 2 independent experiment. Graphs show Mean ± S.D. One-way ANOVA used. ** p=0.0039 (Control vs. Control + Iono), ** p=0.0015 (FMNL1 KD vs. FMNL1 KD + Iono). E: Representative timelapse images of HeLa control, Arf1 KD, & PI4KIIIβ KD cells transfected with FMNL1-GFP (green) & mCherry-TGN46 (magenta) stimulated with 5µM of ionomycin. Arrows indicate vesicles with increasing FMNL1-GFP signal on vesicles. Scale: 5µm. F: Traces showing fold change of FMNL1-TGN46 overlap with 5µM ionomycin stimulation in HeLa control, Arf1 KD, & PI4KIIIβ KD cells transfected with FMNL1-GFP & mCherry-TGN46, normalized to initial intensity prior to stimulation. Data from n = 15 (control), 15 (Arf1 KD), & 13 (PI4K KD) cells from 2 independent experiment. Graph shows mean intensity ± S.E.M. G: Representative fixed-cell images of HeLa Control, Arf1 KD, & PI4KIIIβ KD cells with or without over-expression of GFP-Cdc42-Q61L stained for Tom20 (magenta) Scale: 5µm. H: Quantification of mitochondrial morphology for each parameter set in (G). Quantified for mean area per mitochondria and mean mitochondria per ROI (25µm^2^). Data from n = 25 (Control), 30 (Arf1 KD), 30 (PI4KIIIβ KD), 20 (Control+Cdc42-Q61L), 27 (Arf1 KD+Cdc42-Q61L), 26 (PI4KIIIβ KD+Cdc42-Q61L) from 2 independent experiment. Graphs show Mean ± S.D. One-way ANOVA used. **** p<0.0001.

### FMNL1 mediated actin filaments catalyze the final step in mitochondrial division

We wanted to validate the identity of actin filaments catalyzing the final steps of mitochondrial division. To do this we transfected HeLa cells with ER-Sec61β, mito-BFP and mCherry-Ftractin to visualize ER, mitochondria and actin filaments during unstimulated mitochondrial division events. Both ER and actin filaments were seen to wrap mitochondria prior to mitochondrial division (**Fig 6A**). Approximately 76% of the spontaneous mitochondrial division were positive for actin filaments (**Fig 6A, D**). However, the orientation of the actin signal did not overlap with the ER, suggesting that the actin filaments present at these site were less likely to be INF2-mediated. We then evaluated the few spontaneous division events in FMNL1 KD cells using the same organelle markers. While ER was present (and in a significantly increased abundance) at the mitochondrial division sites, prevalence of actin filaments dropped to only 35% of mitochondrial division in these cells (**Fig 6B, D**). We also carried out similar experiments in INF2-KO cells. The few division events seen in these cells were found positive for both ER and actin filaments, with the latter present in 76% of the mitochondrial division events (**Fig 6C, D**). We also evaluated the kinetics of actin recruitment with respect to FMNL1 appearance at the division sites in both stimulated and unstimulated mitochondrial division. Actin filaments decorated the mitochondrial division sites slightly later than FMNL1 for both unstimulated and stimulated mitochondrial division (**Fig 6E**). Overall, this suggested that actin filaments catalyzing the final steps of mitochondrial division were assembled by FMNL1 and not INF2. To further test the importance of actin filaments in this process, we either overexpressed WT or constitutive active INF2 disease mutant (INF2-R218Q) in control and FMNL1 depleted cells.

**Figure 6.**
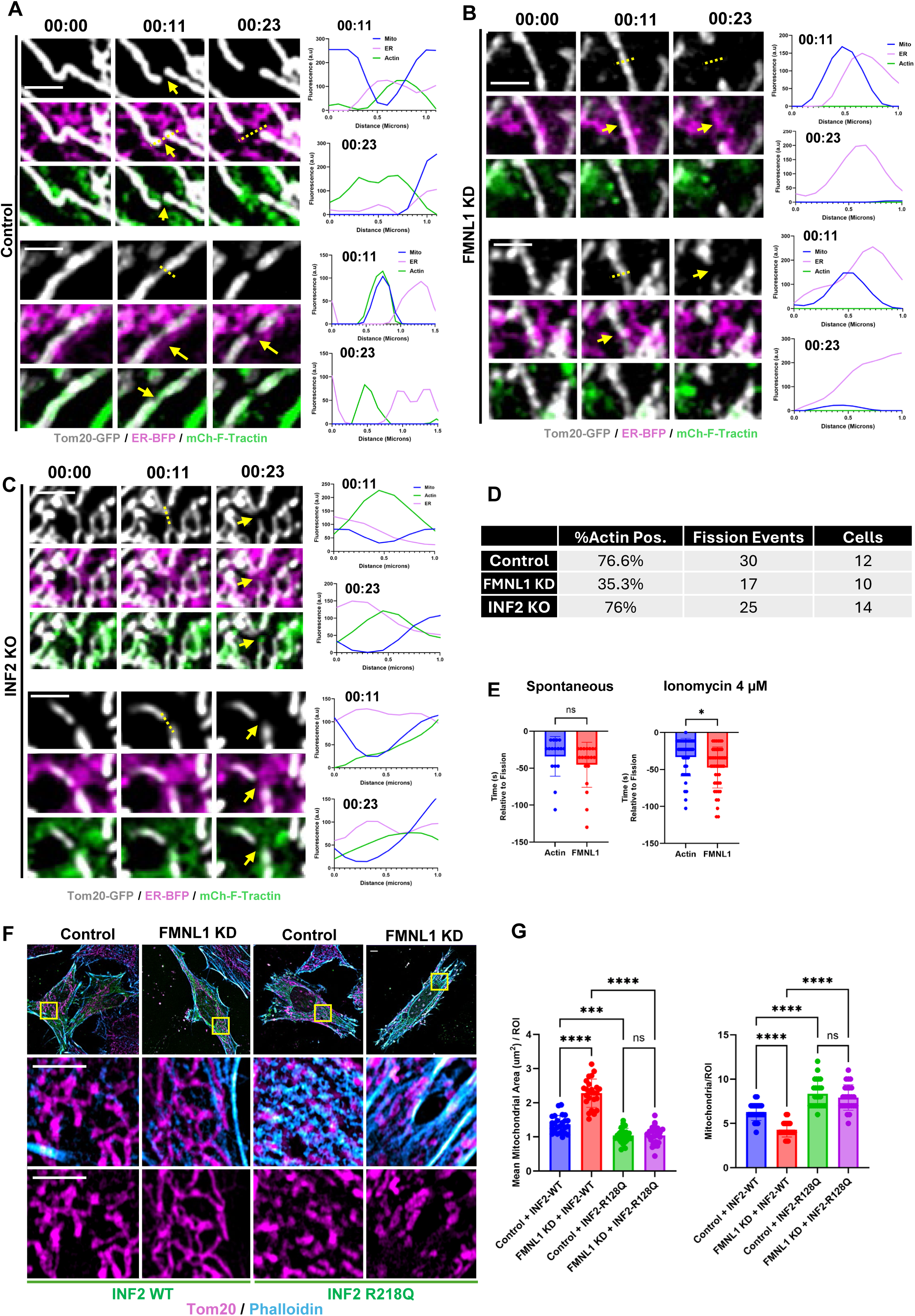
FMNL1 mediated actin filaments catalyze the final step in mitochondrial division. A: (Left) Representative timelapse images of a mitochondrion undergoing division in HeLa control cells transfected with ER-BFP (magenta), Tom20-GFP (gray), mCherry-Ftractin (green). Arrows indicate site of division and dotted line indicates region selected for line scan. (Right) line scan showing fluorescence intensities at indicated time stamps of each fluorophore. B: (Left) Representative timelapse images of a mitochondrion undergoing division in HeLa FMNL1 KD cells transfected with ER-BFP (magenta), Tom20-GFP (gray), mCherry-Ftractin (green). Arrows indicate site of division and dotted line indicates region selected for line scan. (Right) line scan showing fluorescence intensities at indicated time stamps of each fluorophore. C: (Left) Representative timelapse images of a mitochondrion undergoing division in HeLa INF2 KO cells transfected with ER-BFP (magenta), Tom20-GFP (gray), mCherry-Ftractin (green). Arrows indicate site of division and dotted line indicates region selected for line scan. (Right) line scan showing fluorescence intensities at indicated time stamps of each fluorophore. D: Quantification of actin positive mitochondrial division events (shown as % of total division events) from images taken in Fig 6A-C. Data quantified from 12 cells, 30 fission events (control), 10 cells, 17 fission events (FMNL1 KD) and 14 cells, 25 fission events (INF2-KO) across 3-5 independent coverslips in 2 experiments. E: Quantification of arrival times (relative to fission: time 0) of fission factors (Actin and FMNL1) during unstimulated and Ionomycin stimulated division events in HeLa-WT cells. Data from n = 16 (Actin; unstimulated), 19 (FMNL1; unstimulated) division events from 10-12 cells and n = 44 (Actin; stimulated), 47 (FMNL1; stimulated) division events from 12-15 cells across 3 independent experiments Graphs show Mean ± S.D. * p=0.018. F: Representative fixed-cell images of HeLa control & FMNL1 KD cells transfected with either INF2-WT (green) or INF2-R218Q (green) mutant and stained for Tom20 (magenta) and Phalloidin (cyan). Scale: 10µm (main); 5µm (inset) G: Quantification of mitochondrial morphology for each parameter set in (D). Quantified for mean area per mitochondria and mean mitochondria per ROI (25µm^2^). Data from n = 25 (Control+INF2-WT), 24 (FMNL1 KD+INF2-WT), 25 (Control+INF2-R218Q), & 25 (FMNL1 KD+INF2-R218Q) from 2 independent experiment. Graphs show Mean ± S.D. One-way ANOVA used. **** p<0.0001, *** p=0.0009. H: Working model of FMNL1-mediated mitochondrial division.

While overexpression of WT-INF2 did not modify the elongated mitochondrial morphology in FMNL1 KD cells, expression of INF2-R218Q, which results in a constitutive increase in abundance of actin filaments, could significantly fragment mitochondria in FMNL1 KD cells (**Fig 6F, G**). Cumulatively these results suggest that FMNL1 assembled actin filaments catalyze the final steps of mitochondrial division in a Drp1-dependent manner. Loss of FMNL1 can be completely abrogated by expression of a constitutive-active INF2 which can provide actin filaments to fuel and complete the process in the absence of FMNL1.

## Discussion

Collectively we show that FMNL1 (and not FMNL2 or 3) is involved in mitochondrial division. Both acute and chronic depletion of FMNL1 increases mitochondrial length, connectivity and decreases mitochondrial division events in HeLa and U2-OS cells. Furthermore, we show that FMNL1 localizes at mitochondrial division sites being drafted after Drp1recruitment along with TGN vesicles where it is activated specifically by Cdc42. Arf1 and PI(4)KIIIβ operate independent of FMNL1 in regulating mitochondrial division and loss of these factors can be partially overcome by activation of FMNL1. Finally, we show that actin filaments that mediate the final step of mitochondrial scission is polymerized by FMNL1 as the few division events in FMNL1 depleted cells are independent of actin filament assembly but positive for the presence of ER (and therefore INF2). Using these results, we drafted a timeline of the late-stage events of mitochondrial division where FMNL1 is recruited to TGN vesicles (carrying Arf1 and PI(4)KIIIβ) from cytosol (dependent of increased calcium) and is recruited to mitochondrial division sites after Drp1 recruitment. Cdc42 activated FMNL1 at these sites polymerize actin filaments to promote Drp1 GTPase activity which results in mitochondrial division.

Calcium serves as a master regulator of mitochondrial division, stimulating several steps in the process. (1) INF2 is activated in a calcium-calmodulin dependent manner (Lee et al., 2026) that polymerizes actin filaments to promote ERMC and mitochondrial calcium uptake (Chakrabarti et al., 2018). (2) Increased matrix calcium have been shown to induce “constrictions” (Chakrabarti et al., 2018; Cho et al., 2017) or “pearling” (Landoni et al., 2026) which serves as a pre-requisite for IMM division (Chakrabarti et al., 2018) and proper distribution of mtDNA (Landoni et al., 2026). (3) Sustained cytosolic calcium spike also activates phosphatase calcineurin to promote translocation of Drp1 to mitochondria (Cereghetti et al., 2008). (4) We show that increase in cytosolic calcium stimulates translocation of FMNL1 from cytosol to TGN vesicles. FMNL1 myristoylation at its N-term is imperative for its membrane localization and downstream activity (Han et al., 2009). Increase in calcium ions has been shown to cause a myristoyl switch in several proteins that induces expulsion of the myristoyl group resulting in a prompt and reversible interaction with lipid membranes (Ames et al., 1997; Seykora et al., 1996). A similar mechanism might work to stimulate recruitment of FMNL1 to the TGN membranes following ionomycin stimulation. (5) Furthermore, we show that increase in cytosolic calcium results in FMNL1 independent PI4-P formation at the TGN vesicles required for mitochondrial division.

While PI(4)KIIIβ has not been shown to be directly regulated by calcium, increase in PI4-P following ionomycin stimulation might be through an indirect effect on the kinase or its phosphatase Sac1. Indeed, neuron localized synaptojanin is activated through an increase in calcium ions have been shown to promote an increase of PI4-P (Guo et al., 2025).

Small GTPase Cdc42 have emerged as an important mediator of mitochondrial division in several studies. Fasting-induced mitochondrial fission utilizes mTORC2 induced phosphorylated NDRG1 that promote Cdc42 activated actin filaments at fission sites (Martinez-Lopez et al., 2023). More recently FGD1 dependent Cdc42 activation has been proposed to mediate actin assembly and fragmentation of mitochondrial network in Caenorhabditis elegans (Zhao et al., 2026). While these studies outlined the role of Cdc42, our study identifies FMNL1 as the actin nucleator downstream Cdc42 that drives actin-assembly to promote mitochondrial division. Our results indicate that activate Cdc42 can localize both on mitochondria and TGN vesicles and drive mitochondrial fragmentation in FMNL1 and Drp1 dependent manner. Surprisingly, Cdc42 activation can cause mitochondrial fragmentation in INF2 depleted cells, suggesting that actin assembly through FMNL1 can stimulate division in the absence of INF2. Distinct GEFs like FGD1(Zhao et al., 2026) and ARHGEF10 (Martinez-Lopez et al., 2023) might be required to activate Cdc42 in a context dependent manner to stimulate mitochondrial division.

Actin filaments play distinct roles throughout the process of mitochondrial division. INF2 polymerized filaments initially potentiate IP3R mediated calcium release and additionally regulate proper IP3R cluster positioning to mediate ERMC formation and calcium transfer (Zavala et al., 2026). While actin assembly does not require additional factors, mitochondrial calcium transfer requires the presence of non-muscle Myosin II family (Kage et al., 2022; Korobova et al., 2014). Actin filaments were also shown to not only induce productive Drp1 oligomerization at the division site (Ji et al., 2015), but also to promote Drp1 GTP hydrolysis (Hatch et al., 2016). However, the kinetics of INF2-mediated actin assembly and Drp1 oligomerization at division site presented a disconnect (Ji et al., 2015) that raised questions regarding the identity of the actin filaments that promote Drp1 GTP hydrolysis during mitochondrial division. Our results indicate that the final steps of mitochondrial division is catalyzed by FMNL1 polymerized actin filaments. Over-expression of active Cdc42 caused mitochondrial fragmentation in control, INF2, Arf1 and PI(4)KIIIβ depleted cells but not in FMNL1 and Drp1 knock down cells suggesting that the actin filaments generated downstream of Cdc42 were polymerized by FMNL1 and operated in a Drp1 dependent manner. In addition to its nucleation function, FMNL1 can also bundle actin filaments due to the presence of positively charged FH2 domain and dissociable FH2 dimers (Harris et al., 2006) which might be able to provide additional force further supplementing the constriction of OMM through Drp1 GTP hydrolysis.

## Acknowledgments

We thank Tim Ryan (Weill Cornell Medicine) for sharing the ER-GCaMP150 calcium sensor, Gyorgy Hajnoczky (Thomas Jefferson University), Henry Higgs (Geisel School of Medicine at Dartmouth), Frieda Kage (Geisel School of Medicine at Dartmouth) and Roberto Dominguez (University of Pennsylvania) for their critical advice and reagents, and to Caal Kworf for his help in timely completion of the experiments. We would also like to thank Zuzana Nichtova and Nina Petrova from Mitocare TEM Core facility for their support in TEM studies.

## Author Contributions

Conceptualization: BT, AG, RC

Methodology: BT, AG, RC

Investigation: BT, AG, MRZ, RC

Visualization: BT, AG, MRZ, RC

Funding acquisition: RC

Project administration: RC

Supervision: RC

Writing – original draft: BT, RC

Writing – review & editing: BT, RC

## Supplementary Materials

### Materials and Methods

#### Cell culture

Human cervical cancer cells (HeLa) and human osteosarcoma cells (U2-OS) were obtained from American Type Culture Collection (ATCC). U2-OS-GFP-Drp1 knock in cell line was a kind gift from Dr. Henry N. Higgs (Geisel School of Medicine at Dartmouth). Cells were grown in regular DMEM (Gibco 11-995-065) supplemented with 10% fetal bovine serum at 37°C with 5% CO2. Cell lines were tested every 3 months for mycoplasma contamination using Universal Mycoplasma detection kit (ATCC, 30-1012K) or MycoAlert Plus Mycoplasma Detection Ki (Lonza, LT07-701).

#### Generation of FMNL1-KO HeLa cell line

HeLa cells were genome-edited for deletion of FMNL1 gene using the CRISPR/Cas9 technology. CRISPR-gRNA for FMNL1 was as follows: 5’-CACCGAGCGCGGCGTCCCATCGGGG-3 that was designed to target a specific genomic region in exon 1. Selected gRNAs were cloned into pSpCas9(BB)-2A-Puro (Addgene #48139), sequence-verified, and 1 µg of respective plasmids were transfected using Lipofectamine2000. Next day, transfected cells were replated into medium containing 2.5 µg/ml puromycin (Gibco #A1113802) and selected for 24 hours. After 24 hours puromycin containing media along with dead cells were replaced with normal DMEM and the adherent cells were allowed to become confluent. The cells were then diluted to ∼0.5 cells per well and seeded in a 96 well plate to obtain single cell derived clones. Single cell clones were expanded and screened for the absence of FMNL1 protein expression by western blotting using specific antibodies.

#### Plasmids

ER GCaMP6 150 (Kd= 150 µM) was a kind gift from Timothy Ryan (Weil Cornell Medicine, NYC, NY), Mito GCaMPf6 (Kd=0.2 µM), Drp1-YFP, mCherry-OMP-25 and mito-mDendra2 was a kind gift from Gyorgy Hajnoczky (Thomas Jefferson University, Philadelphia, USA). GFP-Ftractin, mitoBFP, mitoGFP, mcherry-Ftractin, GFP-INF2-noncaax, GFP-INF2-non-caax R218Q, eGFP-Tom20, ER-RFP, ER-BFP, GFP-Cdc42-L61, GFP-Rac1-L61, Cdc42-G12V and GFP-Sec61β were a kind gift from Henry Higgs (Geisel School of Medicine, Dartmouth). FMNL1-GFP was a kind gift from Klemens Rottner (Helmholtz Centre for Infectious Research). mCh-Drp1 was a gift from Gia Voeltz (Addgene plasmid # 49152; http://n2t.net/addgene:49152; RRID:Addgene_49152). mCherry-TGNP-N-10 was a gift from Michael Davidson (Addgene plasmid # 55145; http://n2t.net/addgene:55145; RRID:Addgene_55145). Arf1-GFP was a gift from Paul Melancon (Addgene plasmid # 39554; http://n2t.net/addgene:39554; RRID:Addgene_39554). GFP-P4M-SidMx2 was a gift from Tamas Balla (Addgene plasmid #51472; http://n2t.net/addgene:51472; RRID:Addgene_51472)

For all experiments, the following amounts of DNA were transfected per 6-well (individually or combined for co-transfection): 200 ng of mito-GCaMPf6, Drp1-YFP, mCherry OMP25, mito-mDendra2, mitoBFP, mitoGFP, mCherry-Drp1, mCherry-TGNP-N-10, Arf1-GFP, Tom20 GFP, FMNL1-GFP and GFP-P4M-SidMx2. 400 ng of mcherry-Ftractin, GFP-Cdc42-L61, GFP-Rac1-L61 and GFP-Ftractin; 500 ng of ER GCaMP6 150, ER-RFP, ER-BFP and GFP-Sec61β; 1000 ng of GFP-INF2-noncaax, GFP-INF2-noncaax R218Q.

#### Plasmid transfections

Cells were seeded at 4×10^5^ cells per well in a 35 mm dish at ∼16h before transfection. Plasmid transfections were made in OPTI-MEM media (Gibco, 31985062) containing 2.5 µl of Lipofectamine 2000 (Invitrogen, 11668) following the manufacturer’s protocol. After 4-6 h of transfection, cells were trypsinized and re-plated onto glass bottomed dishes (MatTek Corporation, P35G-1.5-14-C) at ∼1×10^5^ cells per well (for live cell imaging) or onto coverslips (25/10 mm) for fixed cell analysis. Cells were imaged or fixed ∼16–24 h after transfection.

#### siRNA induced knockdown

Cells were seeded at 1×10^5^ cells per well in a 35 mm dish ∼ 16 h before transfection. 63 pg of siRNA was added to OPTI-MEM (Gibco, 31985062) with 2 µl RNAimax (Invitrogen, 13778100), incubated at RT for 20 minutes and added to cells contained in 1 ml of pre-warmed DMEM media. 72-96 hours later cells were transfected with respective reporter constructs and prepared for live/fixed cell analysis. Following siRNA oligos were used:

INF2-: 5’-GGAUCAACCUGGAGAUCAUCCGC-3’ (Custom made)

FMNL1-5’-GTGGTACATTCGGTGGATCATGTTCTCCACCGAAT-3’(IDT; hs.Ri.FMNL1.13.5)

FMNL2-5’-CATGATGCAGTTTAGTAA-3’ (IDT; hs.Ri.FMNL2.13.1)

FMNL3-5’-GCATCAAGGAGACATATGA-3’ (Ambion; Cat#s40551)

Drp1-5’-GCCAGCTAGATATTAACAACAAGAA-3’ (Custom made)

Arf1-5’-CAAUUCUGCAUGGUCACAGUAGAGA-3’ (IDT; hs.Ri.ARF1.13.1)

Arf1-5’-UCUCUACUGUGACCAUGCAGA AUUGAU-3’ (IDT; hs.Ri.ARF1.13.2)

PI4KIIIβ- 5’-CCCCAACUGUUUGUUAGUAUUAUTA-3’ (IDT; hs.Ri.PI4KB.13.1)

PI4KIIIβ- 5’-AUCAUGUGGUAACUGCGAGAGCUTG-3’ (IDT; hs.Ri.PI4KB.13.4)

#### Drug treatments

To evaluate the agonist-induced ER calcium release and mitochondrial calcium uptake 1 uM histamine (H7125; Sigma-Aldrich) was used. Agonist was perfused manually at the fifth frame (∼25 sec) during the imaging and not replaced during the entire protocol (5 min). Mitochondrial fragmentation was induced by 1 µM staurosporine (19-123-M; Sigma Aldrich) or 10µM BAPTA-AM (A1076; Sigma-Aldrich). Ionomycin (I3909; Sigma-Aldrich) was used at 4 µM to induce mitochondrial fission in live cell experiments. Quant-iT™ PicoGreen™ dsDNA Assay Kits (P7589; Thermo Fischer Scientific) was used to stain mtDNA in live cells.

#### Immunofluorescence

Cells (transfected or un-transfected) were plated onto coverslips ∼16 h prior to fixation and staining. Cells were then fixed with either 1% Glutaraldehyde (for mitochondrial length evaluation) or pre-warmed 4 % PFA (for all other experiments). In case of Glutaraldehyde fixation, post-fixation cells were washed with PBS and incubated with freshly made Sodium borohydride (1 mg/ml) 15 mins (X3). Cells were then washed with PBS, permeabilized with 0.25% TX-100 (10 minutes). For PFA fixed samples, cells were directly permeabilized with 0.1% Triton X-100 for 1 min and washed with PBS three times. In all conditions cells were blocked with 10% FBS in PBS for ∼60 min. Primary antibody was diluted in 1% FBS/PBS and coverslips were incubated on a drop of antibody solution on parafilm in a wet chamber for 1 h. Cells were then washed 6 times with 1X PBS and incubated with appropriate secondary antibody (prepared in 1% FBS/PBS) for 1 hour. Coverslips were again washed 6 times with 1X PBS and mounted on glass slides using ProLong Gold antifade mounting media (Invitrogen #P36930).

<u>Primary antibodies:</u>

anti-Tom 20 (Abcam ab78547, used at 1:400)

anti-Dlp1 (BD Transductions laboratories 611113, used at 1:200)

<u>Secondary antibodies:</u>

Goat anti-Mouse IgG (H+L) Cross-Adsorbed Secondary Antibody, Alexa Fluor™ 647 (Invitrogen #A21235) used at 1:1000; Goat anti-Rabbit IgG (H+L) Cross-Adsorbed Secondary Antibody, Alexa Fluor™ 488 (Invitrogen #A11008) used at 1:1000

#### Proximity Ligation Assay (PLA)

PLA (Duolink™ PLA technology, *In Situ* Starter Kit Mouse/Rabbit, MilliporeSigma, cat# DUO92101) was used to evaluate the interaction between VAPB/Tom20. Briefly, cells were seeded onto 12 mm coverslips for 16 hours and then were fixed with prewarmed 4% PFA for 20 min at RT. Cells were washed with PBS (X 3 times) and permeabilized using 0.1% Triton-X-100 for 1 min. The cells were then incubated with Duolink blocking buffer for 1 hour and incubated with primary antibodies (diluted in Duolink antibody diluent) for 2 hours at RT. Cells were then washed with 5% BSA (15 min x 3) and incubated with Duolink secondary antibodies for 1 hr at 37C. Cells were then washed with Duolink wash buffer A (5 min x 3) and incubated with Duolink ligase for 30 min at 37C. Cells were further washed with Duolink wash buffer A (2 min x 3) and incubated with Duolink polymerase for 90 min at 37 C. The cells were finally washed with Duolink wash buffer B (10 min x 3) and mounted on glass slides using prolong gold antifade reagent. Images were acquired within 24-48 hours. For VAP-B/Tom20 PLA staining, the Alexa fluor 488 Phalloidin was added in the blocking buffer to mark actin filaments.

<u>Antibodies and stains used:</u>

anti-VAP-B (Proteintech 66191-1-Ig used at 1:400)

and anti-Tom20 (Abcam ab78547, used at 1:400)

Alexa fluor 488 Phalloidin (Invitrogen, A12379, used at 1:500)

#### Lysate preparation and Western Blot

To evaluate protein abundance cells from a 35mm dish were lysed using 1% SDS, collected and heated at 95C for 5 min. To run them on SDS-PAGE, samples were mixed with 2xDB (50mM Tris-HCl, pH 6.8, 2mM EDTA, 20% glycerol, 0.8% SDS, 0.02% Bromophenol Blue, 1M NaCl, 4M urea) and separated in a 4-20% precast gradient gels, transferred to a polyvinylidene fluoride membrane (EMD Millipore, IPFL00010). The membrane was blocked with TBS-T (20 mM Tris-HCl, pH 7.6, 136 mM NaCl, 0.1% Tween20) containing 3% BSA (VWR Life Science, VWRV0332) for 1h, then incubated with primary antibody solution at 4°C overnight or 2 hours at RT. After washing with TBS-T, the membrane was incubated with requisite secondary antibody for 1h at room-temperature and developed on a Chemi-doc system using appropriate reagents.

<u>Primary Antibodies used:</u>

anti-INF2 antibody (Proteintech, 20466-1-AP, used at 1:1000)

anti-β-Actin antibody (CST, #3700S, used at 1:5000; SIGMA, A2228 used at 1:10000)

anti-FMNL1 antibody (Proteintech, 27834-1-AP, used at 1:3000 RT/2 hrs)

anti-FMNL2 antibody (Abcam, ab75963, used at 1:2000)

anti-FMNL3 antibody (Abcam, ab222797, used at 1:2000)

anti-Drp1 antibody (CST, 8570S, used at 1:1000)

anti-OPA1 antibody (BD Transduction laboratories, 612607, used at 1:2500)

anti-Mfn2 antibody (CST, 9482, used at 1:1000)

anti-MFF antibody (Abcam abEPR7360, used at 1:1000)

anti-PTPIP51 (Proteintech, 20641-1-AP, used at 1:1000 at RT/ 2hrs)

<u>Secondary Antibodies used:</u>

anti-Rabbit-HRP antibody (Jackson Immuno Research, 111-035-144, used at 1:10000)

anti-Mouse-HRP (Jackson Immuno Research, 715-035-150, used at 1:5000)

anti-rabbit IgG (H+L) antibody conjugated to IRDye 800CW (LICORBio, 926-32211, used at 1:10000)

anti-Mouse IgG (H+L) antibody conjugated to IRDye 680LT (LICORBio, 926-68020, used at 1:10000)

#### Immunoprecipitation using GFP-Trap agarose beads

Briefly, HEK-293 cells were seeded into a 10 cm dish and were transfected with FMNL1-GFP with or without Cdc42-G12V construct. After 6h of transfection, media was replaced with fresh DMEM. Next day, media was removed, washed with PBS and lysates was prepare using immunoprecipitation buffer (IPB) (IPB: 140 mM KCl, 50 mM Tris-HCl (pH 7.4), 2 mM MgCl_2_) with protease and phosphatase inhibitors containing 1% Triton-X100, Lat A (2 µM) and Phalloidin (10 µM). Cells were lysed on a rotator for 1 h at 4C, then centrifugated for 15 min at 15000 rpm at 4C to collect the supernatant. ∼10% of the volume was used for the *Input*, and the rest was added to the conditioned GFP-trap-beads. After 2 h mixing on a rotator at 4C, the beads were collected via centrifugation, washed with IPB (X 3) and reconstituted in sample buffer, heated at 95C for 5min, and evaluated through SDS-PAGE.

#### Microscopy

Live-cell images pertaining to Figures 1G, 1H, 3A-H, 4F-H, 5A-D, 5G-H, 6A-B, S2A-F, S2I-J, S4H-J and S5E were acquired using a Nikon A1 on a Nikon Ti-E base and equipped with an iXon Ultra 888 EMCCD camera. A solid-state 405 smart diode 100-mW laser, solid-state 488 OPSL smart laser 50-mW laser, solid-state 560 OPSL smart laser 50-mW laser, and solid-state 637 OPSL smart laser 140-mW laser were used (objective: 20X, 1.45 NA CFI Plan Apo; Nikon). Images were acquired using Nikon Elements v5.02 and quantified using Image J.

Live and Fixed cells pertaining to Figures 1A-F, 2A-B, 2G-J, 4A-E, 5E-F, 6C-D, 7B-D, S1B-C, S1E-F, S2G-H, S2K-L, S3B-H, S4A-G and S5A-D were performed on LSM 880 equipped with 63×/1.4 NA plan Apochromat oil objective using the Airyscan detectors (Carl Zeiss Microscopy). The Airyscan uses a 32-channel array of GaAsP detectors configured as 0.2 airy units per channel to collect data that is subsequently processed using Zen2 software. Cells were imaged with the 405-nm laser and 450/30 filter for BFP, 488 nm and 525/30 for GFP, 561 nm and 595/25 for RFP and 640 nm laser and 650 nm LP for Far-red. Images were quantified using Image J.

#### Calcium measurements

Cells expressing either ER or mitochondrial calcium sensors were washed twice with PBS1X and perfused with 0.25 % BSA ECM buffer, calcium-free (ECM: 121 mM NaCl, 5 mM NaHCO_3_, 10 mM Na-HEPES, 4.6 mM KCl, 1.2 mM KH_2_PO_4_, 1.2 mM MgSO_4_, 10 mM glucose, pH7.4) and transferred to the microscope equipped with a heated (37C) stage-top. Cells were recorded for 25 seconds at 5s/frame, stimulated with agonists and imaged for another 5 minutes at 5s/frame. Images were quantified using ImageJ where values were expressed as fold change F/F0 where F0 is the mean of resting fluorescence intensity before stimulus addition.

#### Actin burst

To measure actin burst, HeLa control and FMNL1 KD cells were seeded onto coverslips and allowed to grow for 16-24 hours. Cells were then stimulated with 4 uM ionomycin for 45 seconds, fixed with 4% pre-warmed PFA for 20 min at RT, washed with PBS (X3), permeabilized with 0.1% Triton X-100 for 1 min at RT, washed with PBS (X3) and stained with phalloidin-647 for 30 minutes at RT. The cells were then washed with PBS and mounted on slides using Prolong gold to be imaged.

#### Transmission Electron Microscopy

HeLa cells (control and FMNL1 KD) for TEM analysis were fixed using 2% glutaraldehyde, then with 1% OsO_4_ and stained with 0.5% uranyl acetate, pelleted in 2% agarose (Sigma– Aldrich, Type IX ultralow gelling temperature), dehydrated in a dilution series of acetone/ water, and embedded in Spurr’s resin (Electron Microscopy Sciences). The sections were examined with a JEOL JEM1010 TEM fitted with a side-mounted AMT XR- 50 5Mpx CCD camera or an FEI Tecnai 12 TEM fitted with a bottom-mounted AMT XR-111 10.5 Mpx CCD camera. Images were taken with or ×3200–4400 (FEI) direct magnification. Interfaces between ER and mitochondria were analyzed using a custom ImageJ script: https://sites.imagej.net/MitoCare/ as described previously.

#### Image analysis and quantification

Image J (NIH) was used to analyze and process all images

##### Quantification of calcium transients

Cells transfected with the indicated calcium probes as described above were plated on MatTek dishes for live-cell imaging. Cells were imaged using a 20X water objective on a heated stage at a single confocal slice in the medial region, approximately 2 μm above the basal surface, to avoid stress fibers. Mean fluorescence intensity was calculated for each cell using the ImageJ plug-in “Time Series Analyzer V3.” after background subtraction (taken from extra-cellular 2-3 ROI in the field). Fluorescence values for each time point after drug treatment were normalized to the mean baseline fluorescence (first five frames) and plotted as intensity fold change over time in seconds. Average fluorescence intensities for mitochondrial calcium and ER calcium were determined by outlining the respective organelle networks. Data were plotted as intensity– time curves each data point using Microsoft Excel.

##### Quantification of Drp1-Mito overlap, or TGN-Mito overlap or FMNL1-TGN overlap

Z-stacked images (using Airyscan) from Control and FMNL1 KD cells either stained or expressing appropriate markers were assessed using Image J. We thresholded and extracted the co-localized pixels between the two channels using the Colocalization ImageJ plugin with the following parameters: ratio 50% (0–100%); threshold channel 1: 30 (0–255); threshold channel 2: 30 (0–255); display value: 255 (0–255). Overlapping pixels between Drp1 and mitochondria were normalized to total mitochondrial network. Overlapping pixels between TGN and mitochondria were normalized to the total mitochondrial network. Overlapping pixels between FMNL1 and TGN were normalized to total TGN network. Similar analyses were also performed for Arf1-TGN colocalization and PI4-P-TGN co-localization from live cell images.

##### Quantification of mitochondrial length

Evaluation of mitochondrial length was performed as follows: Briefly maximum intensity projections were generated from z-stacks acquired and background subtracted using FIJI (rolling ball 20.0). A 225-µm^2^ area of resolvable mitochondria was selected (one ROI/cell in the spread region of the cell) and analyzed using “Analyze Particle” plugin in FIJI to obtain number of mitochondrial fragments and the area of each fragment per ROI. Graphs were plotted for number of mitochondria/ROI and Area/mitochondria. Lesser mitochondria occupying bigger area was deemed as elongated mitochondria.

##### Quantification of mitochondrial division rate

Mitochondrial division rate was calculated as follows: Suitable ROIs were selected for analysis based on whether individual mitochondria were resolvable and did not leave the focal plane. Files of these ROIs were assembled, coded and scrambled by one investigator, and analyzed for division by a second investigator in a blinded manner as to the treatment condition. The second investigator scanned the ROIs frame by frame manually for division events and determined mitochondrial length within the ROI using ImageJ. The results were presented as number of fission events/min/1000 um^2^ of mitochondria.

##### Quantification of PLA assays

z- projected images were background-subtracted using “rolling ball radius” plugin of Image J with a value of 20.0 and phalloidin (actin) channel was used to get the cellular area. Channel containing the PLA dots were converted to 8-bit and binarized. “Analyze particles” plugin was used to record total number of dots and presented as number of dots per 100 um^2^ of cellular area. All images collected were quantified.

#### Statistical analysis

All statical analysis and p-values were determined using GraphPad Prism 10. To determine *p*- values, either an unpaired Student’s *t*-test was performed between the two groups of data, comparing full datasets as stated in the Figure legends. To compare across more than two conditions, multiple comparisons (unpaired) and one-way or two-ways ANOVA were performed in GraphPad Prism 10. Data are expressed as Mean±SD.

### Supplementary Figures

**Supplementary Figure 1.**
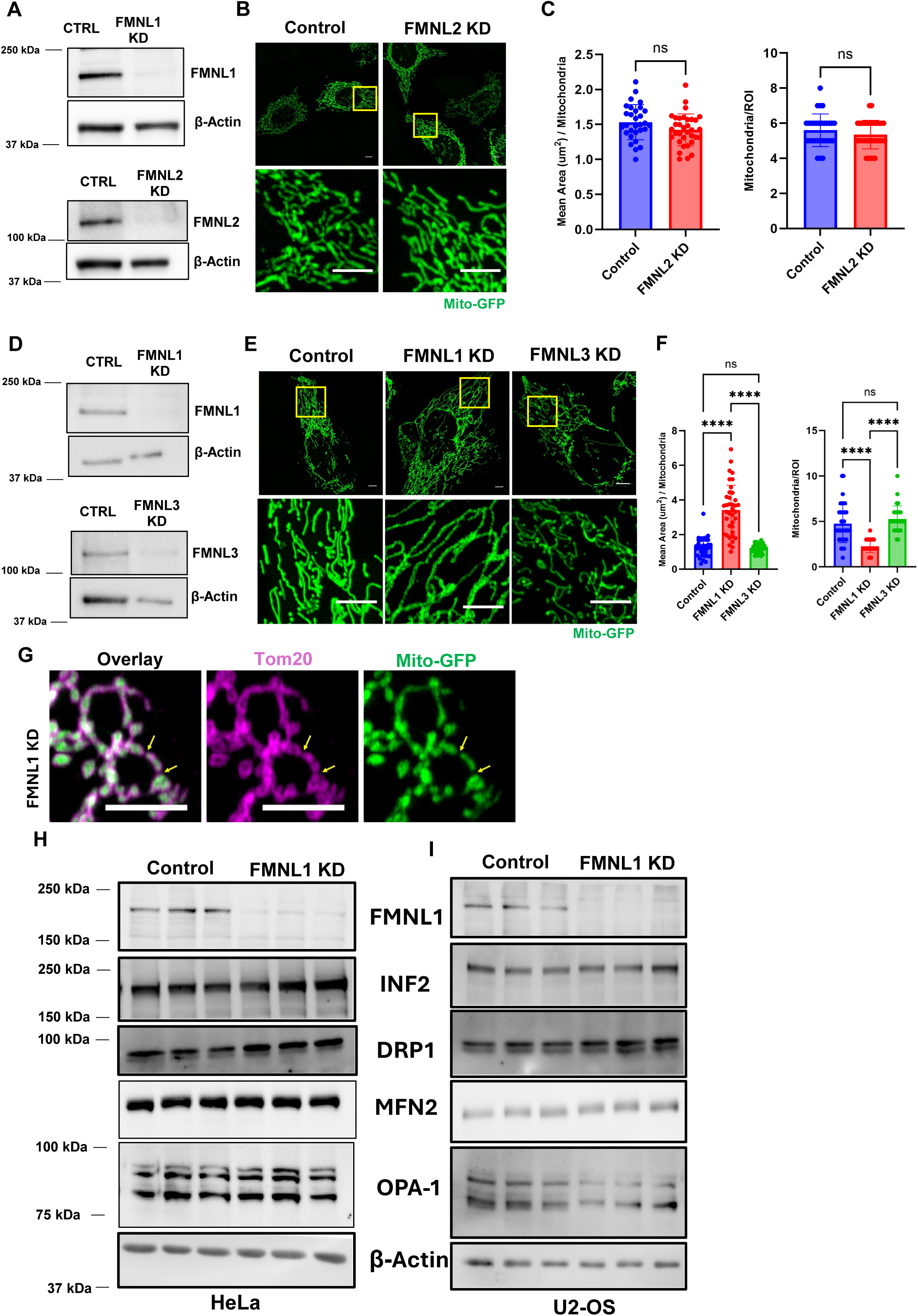
A: (Top) Western blot to evaluate FMNL1 expression in HeLa control & FMNL1 KD cells, β- actin used as loading control. (Bottom) Western blot to evaluate FMNL2 expression in HeLa control & FMNL2 KD cells, β-actin used as loading control. B: Representative fixed-cell images of HeLa control & FMNL2 KD cells transfected with mitoGFP. Boxes indicate regions shown below. Scale: 5µm. C: Quantification of mitochondrial morphology for each condition in (B). Quantified for mean area per mitochondria (left) and mean mitochondria per ROI (25µm^2^) (right). Data from n = 30 (control) & 30 (FMNL2 KD) cells from 3 independent experiment. Graphs show Mean ± S.D. Unpaired student’s t-test used. D: (Top) Western blot to evaluate FMNL1 expression in U2-OS control & FMNL1 KD cells, β-actin used as loading control. (Bottom) Western blot to evaluate FMNL3 expression in U2-OS control & FMNL3 KD cells, β-actin used as loading control. E: Representative fixed-cell images of HeLa control, FMNL1 KD, & FMNL3 KD cells transfected with mitoGFP. Boxes indicate regions shown below. Scale: 5µm. F: Quantification of mitochondrial morphology for each condition in (B). Quantified for mean area per mitochondria (left) and mean mitochondria per ROI (25µm^2^) (right). Data from n = 37 (control), 38 (FMNL1 KD), 30 (FMNL3 KD) cells from 2 independent experiments. Graphs show Mean ± S.D. One-way ANOVA used. **** p<0.0001. G: Representative fixed-cell images of HeLa FMNL1 KD cells transfected with mitoGFP (green) & stained with Tom20 (magenta) following Ionomycin treatment. Arrows indicate regions of interest. Scale: 5µm. H: Western blot to evaluate expression of FMNL1 & known fission/fusion factors as indicated in HeLa control & FMNL1 cells. β-actin used as loading control. I: Western blot to evaluate expression of FMNL1 & known fission/fusion factors as indicated in U2-OS control & FMNL1 cells. β-actin used as loading control.

**Supplementary Figure 2.**
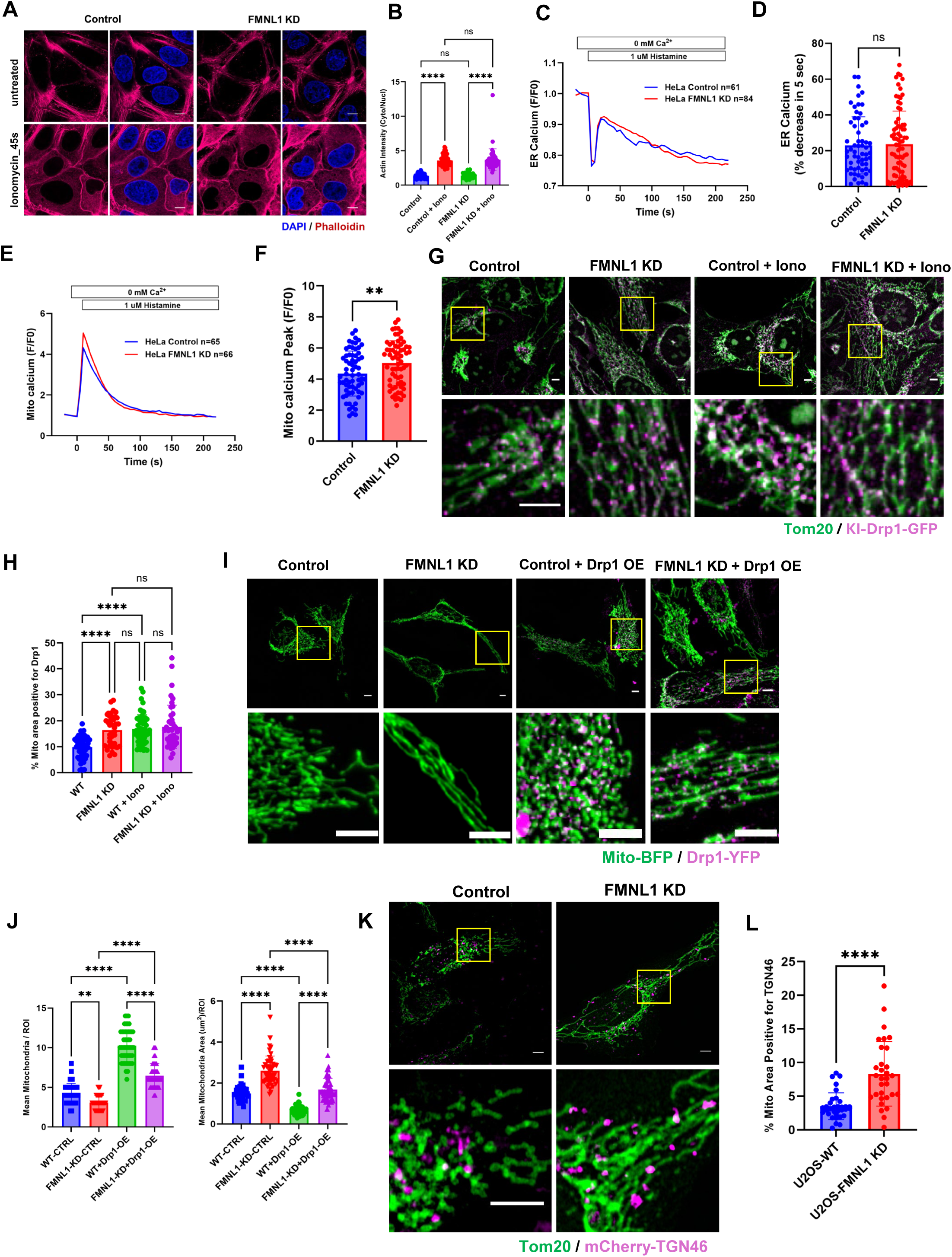
A: Representative fixed-cell images of the medial section of U2-OS control & FMNL1 KD cells stained with Phalloidin (magenta; actin filaments) and DAPI (blue; nucleus) untreated (top) or treated with 4µM ionomycin/ 45 seconds. Scale: 5µm. B: Quantification of (A) measuring actin intensity ratio in the cytosol/nucleus with and without ionomycin stimulation. Data from n = 56 (control), 56 (control+iono), 35 (FMNL1 KD), 48 (FMNL1 KD+iono) cells from 2 independent experiment. Graphs show Mean ± S.D. One-way ANOVA used. ****p<0.0001. C: Average traces of ER calcium transients in HeLa control & FMNL1 KD cells transfected with ER-GCaMP6-150 probe (Kd=150 µM), stimulated with histamine (1µM). D: Quantification of ER calcium expressed as % decrease in 5 sec from traces shown in C. Data from n = 60 (control) & 78 (FMNL1 KD) cells from 3 independent experiment. Unpaired student’s t-test used. E: Traces of mitochondrial calcium transients in HeLa control & FMNL1 KD cells transfected with mito-R-GECO probe (Kd=0.5 µM), stimulated with histamine (1µM). F: Quantification of maximal peak of mitochondrial Ca^2+^ uptake as shown in E, expressed as fold change (F/F0). Data from n = 65 (control) & 66 (FMNL1 KD) cells from 3 independent experiment. Unpaired student’s t-test used. ** p=0.0074. G: Representative fixed-cell images of control and FMNL1 KD U2-OS Drp1-GFP knock-in cells showing Tom20 (green) and Drp1 (magenta) in unstimulated and ionomycin stimulated (4 µM/ 5 min) conditions as indicated. Boxes indicate regions seen in lower panels. Scale 5µm. H: Quantifications of % mitochondrial area positive for Drp1 from data in G. Data from n = 51 (Control), 39 (FMNL1 KD), 53 (Control + Iono), and 43 (FMNL1 KD + Iono) cells from 2 independent experiments. Graphs show Mean ± S.D. One-way ANOVA used. **** p<0.0001. I: Representative fixed-cell images of HeLa control & FMNL1 KD cells transfected with mitoBFP (green) and overexpressing Drp1-YFP (right) (magenta). Boxes indicate regions seen in lower panels. Scale 5µm. J: Quantification of mitochondrial morphology for each condition in (I). Quantified for mean area per mitochondria and mean mitochondria per ROI (25µm^2^). Data from n = 43 (control), 50 (FMNL1 KD), 50 (control+Drp1 OE), 46 (FMNL1 KD+ Drp1 OE) cells from 2 independent experiment. Graphs show Mean ± S.D. One-way ANOVA used. **** p<0.0001, ** p=0.001. K: Representative fixed-cell images of U2-OS Control & FMNL1 KD cells transfected with mCherry-TGN46 (magenta) stained for Tom20 (green). Boxes indicate regions seen in lower panels Scale: 5µm. L: Quantifications of % mitochondrial area positive for TGN46. Data from n = 37 (Control), 32 (FMNL1 KD), & 36 (INF2 KD) cells from 2 independent experiments. Graphs show Mean ± S.D. Unpaired student’s t-test used. **** p<0.0001.

**Supplementary Figure 3.**
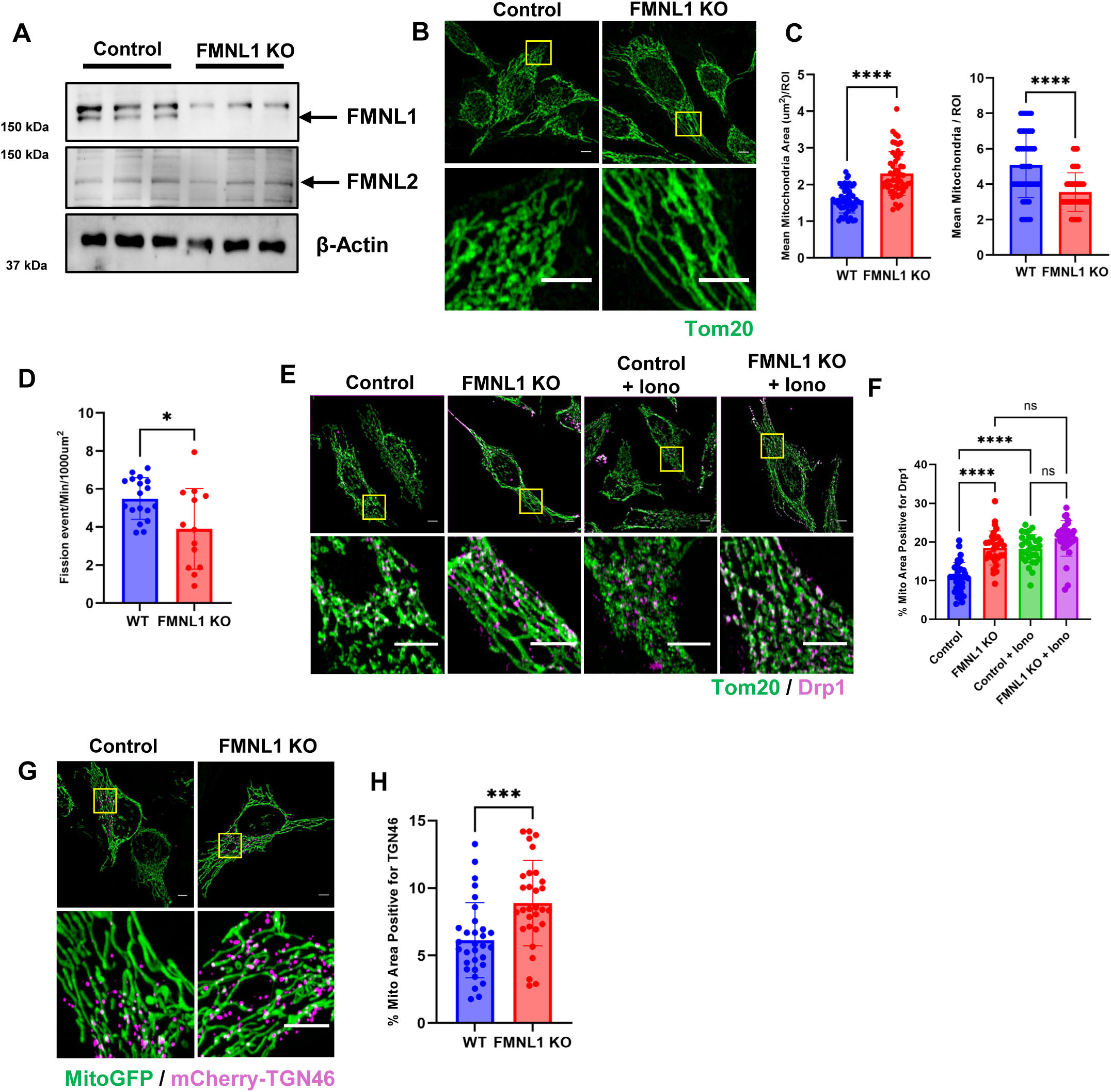
A: Western blot to evaluate expression of FMNL1 & FMNL2 in HeLa control and FMNL1 KO cells. β-actin used as loading control. B: Representative fixed-cell images of HeLa control & FMNL1 KO cells stained with Tom20 (green). Boxes indicate regions shown below. Scale: 5µm. C: Quantification of mitochondrial morphology for each condition in (B). Quantified for mean area per mitochondria (left) and mean mitochondria per ROI (25µm^2^) (right). Data from n = 50 (control) & 52 (FMNL1 KO) cells from 2 independent experiments. Graphs show Mean ± S.D. Unpaired student’s t-test used. **** p<0.0001. D: Quantification of mitochondrial division expressed as events/min/1000µm^2^ in HeLa WT and FMNL1 KO cells. Data from n = 18 (WT) & 13 (FMNL1 KO) cells from 2 independent experiments. Graphs show Mean ± S.D. Unpaired student’s t-test used. * p=0.0107. E: Representative fixed-cell images of HeLa WT & FMNL1 KO stained for Tom20 (green) and Drp1 (magenta) in unstimulated and ionomycin (4µM/ 5 min) stimulated conditions. Boxes indicate regions seen in lower panels. Scale 5µm. F: Quantifications of % mitochondrial area positive for Drp1 from data collected in E. Data from n = 30 (WT), 30 (FMNL1 KD), 30 (Control + Iono), and 30 (FMNL1 KD + Iono) cells from 2 independent experiment. Graphs show Mean ± S.D. One-way ANOVA used. **** p<0.0001. G: Representative fixed-cell images of HeLa WT & FMNL1 KO cells transfected with mitoGFP (green) and mCherry-TGN46 (magenta). Boxes indicate regions seen in lower panels Scale: 5µm. H: Quantifications of % mitochondrial area positive for TGN46 from data collected in G. Data from n = 31 (WT), 30 (FMNL1 KO) cells from 2 independent experiments. Graphs show Mean ± S.D. Unpaired student’s t-test. *** p=0.0006.

**Supplementary Figure 4.**
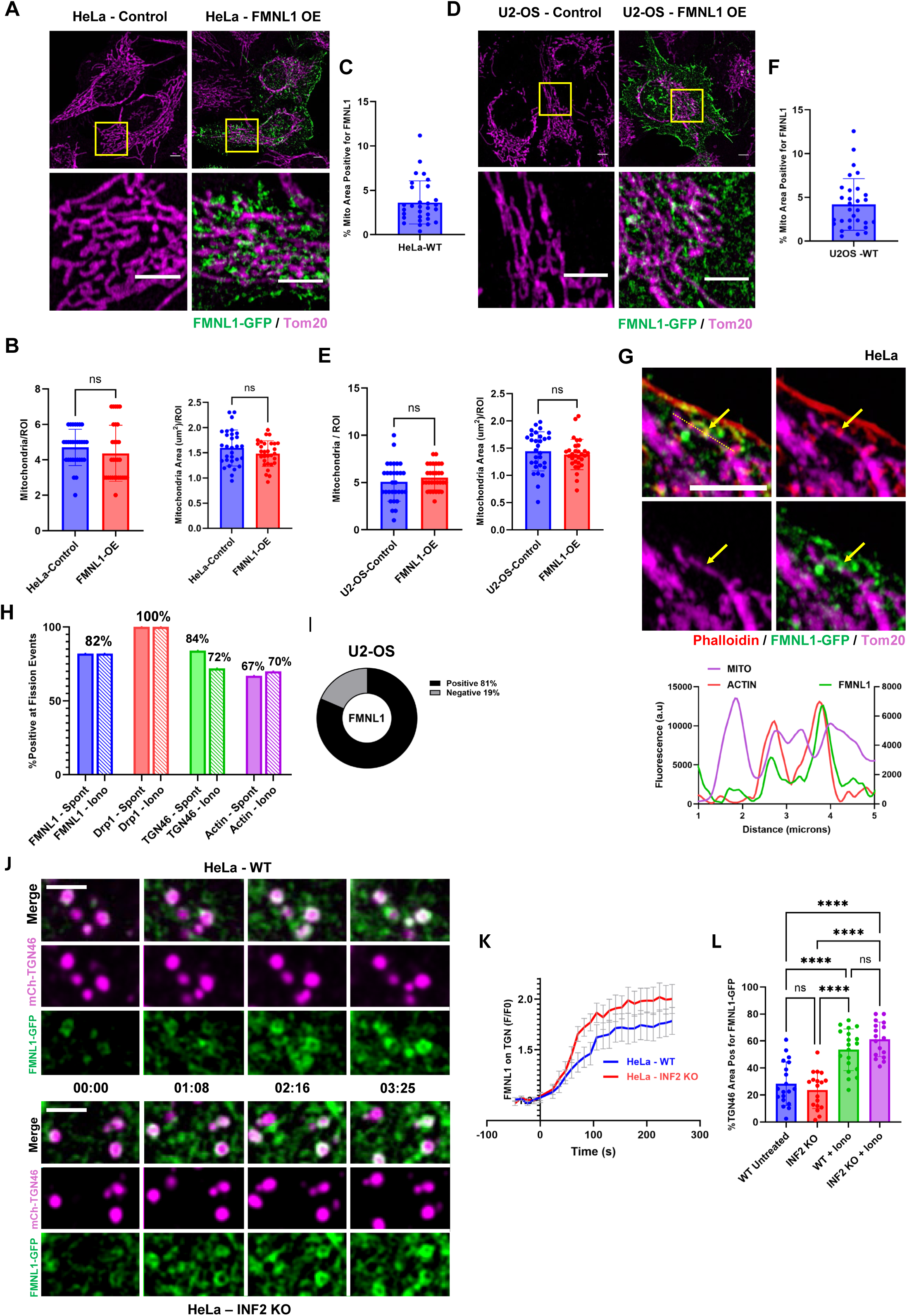
A: Representative fixed-cell images of HeLa cells un-transfected or overexpressing with FMNL1-GFP (green) fixed and stained for Tom20 (magenta). Box indicates region seen in lower panels. Scale: 5µm. B: Quantification of mitochondrial morphology for each condition in (A). Quantified for mean area per mitochondria (right) and mean mitochondria per ROI (25µm^2^) (left). Data from n = 30 (control) & 30 (FMNL1 O.E) cells from 2 independent experiment. Graphs show Mean ± S.D. Unpaired student’s t-test used. C: Quantification of % mitochondrial area positive for FMNL1-GFP in HeLa WT cells. Graphs show Mean ± S.D. Data from n = 30 (WT) cells from 2 independent experiment. D: Representative fixed-cell images of U2-OS cells un-transfected or overexpressing with FMNL1-GFP (green) fixed and stained for Tom20 (magenta). Box indicates region seen in lower panels. Scale: 5µm. E: Quantification of mitochondrial morphology for each condition in (D). Quantified for mean area per mitochondria (right) and mean mitochondria per ROI (25µm^2^) (left). Data from n = 30 (control) & 30 (FMNL1 O.E) cells from 2 independent experiment. Graphs show Mean ± S.D. Unpaired student’s t-test used. F: Quantification of % mitochondrial area positive for FMNL1-GFP in U2-OS WT cells. Graphs show Mean ± S.D. Data from n = 30 (WT) cells from 2 independent experiment. G: Representative fixed-cell image of a mitochondrial constriction in a HeLa WT cell transfected with FMNL1-GFP (green) and stained for Tom20 (magenta) and Phalloidin (red). Arrow indicates point of constriction and dotted line correlates with line scan graph below. H: Quantification of % mitochondrial division events (unstimulated-filled bars; stimulated- dashed bars) positive for FMNL1, Drp1 and TGN46 from images done in Fig 3. I: Quantification of % mitochondrial division events positive for FMNL1-GFP puncta in HeLa WT (top) and U2-OS WT (bottom) cells. Data from n = 106 (HeLa) and 243 (U2-OS) division events from 14 (HeLa) and 26 (U2-OS) cells from 2 independent experiment. J: Representative live-cell timelapse images of HeLa WT (top) and HeLa INF2 KO (bottom) cells transfected with FMNL1-GFP (green) and mCherry-TGN46 (magenta) after stimulation with 5µM ionomycin. Scale: 5µm. K: Traces of FMNL1-TGN46 colocalization (expressed as fold change) in HeLa WT and INF2 KO cells after stimulation of 5µM ionomycin. Error bars shown as S.E.M. Data is from n = 16 (WT) and 15 (INF2 KO) cells across 2 independent experiments. L: Quantification of %TGN area positive for FMNL1-GFP before and after stimulation with 5µM ionomycin in HeLa WT & INF2 KO cells. Data is from n = 19 (WT) and 18 (INF2 KO) cells from 3 independent experiments. Graphs show Mean ± S.D. One-way ANOVA used. **** p<0.0001.

**Supplementary Figure 5.**
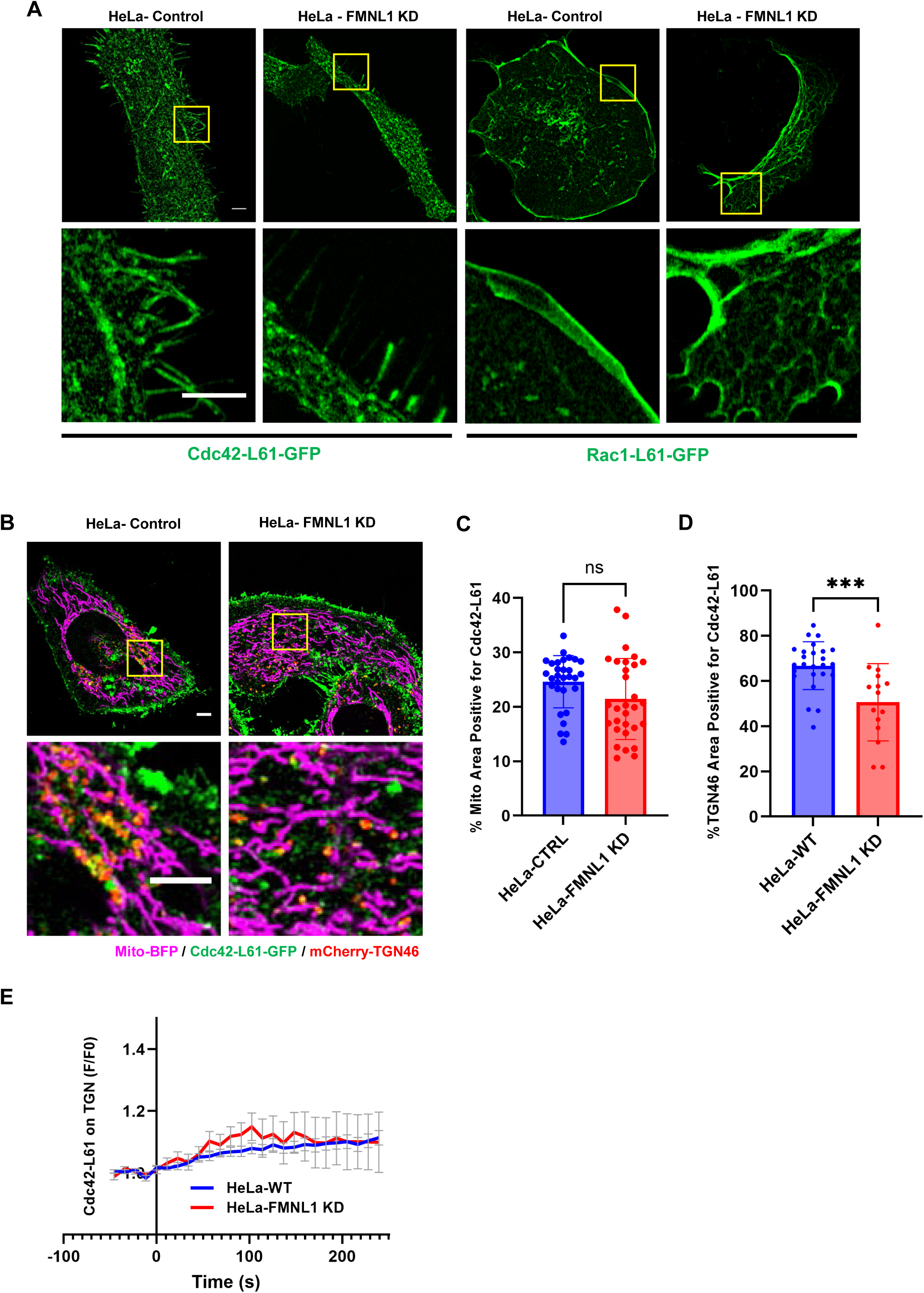
A: Representative fixed-cell images of HeLa control & FMNL1 KD cells transfected with either GFP-Cdc42-Q61L (left) or GFP-Rac1-Q61L (right) (both in green). Boxes indicate regions shown in lower panels. Scale: 5µm. B: Representative live-cell images of HeLa control & FMNL1 KD cells transfected with mitoBFP (magenta), GFP-Cdc41-Q61L (green), and mCherry-TGN46 (red). Boxes indicate regions shown in lower panels. C: Quantification of % mitochondrial area positive for GFP-Cdc42-Q61L puncta in HeLa control and FMNL1 KD cells. Data from n = 30 (control) and 30 (FMNL1 KD) cells from 2 independent experiment. Graphs show Mean ± S.D. Unpaired student’s t-test used. D: Quantification of %TGN46 area positive for GFP-Cdc42-Q61L puncta in HeLa control and FMNL1 KD cells. Data from n = 25 (control) and 15 (FMNL1 KD) cells from 2 independent experiments. Graphs show Mean ± S.D. Unpaired student’s t-test used. *** p=0.0007. E: Traces of Cdc42Q61L-TGN46 colocalization in HeLa WT and FMNL1 KD cells after stimulation of 5µM ionomycin shown as fold change. Error bars shown as S.E.M. Data is from n = 25 (WT) and 15 (FMNL1 KD) cells across 2 independent experiments.

